# Replication-driven HBV cccDNA loss in chimeric mice with humanized livers

**DOI:** 10.1101/2023.12.28.573542

**Authors:** Bai-Hua Zhang, Yuanping Zhou, Ben Tempel, Hongguang Pan, Stephen Horrigan, Laura Luckenbaugh, Fabien Zoulim, Jianming Hu, Yong-Yuan Zhang

**Affiliations:** Virology, HBVtech, Rockville, MD; Department of Hepatology, Nafang Hospital, Southern Medical University, Guangzhou, China; In Celtu, LLC, Rockville, MD; Preclinical Department, Noble Life Sciences, Sykesville, MD; I Centre de Recherche en Cancerologie de Lyon (CRCL), INSERM U1052, CNRS UMR 5286, Université Claude Bernard Lyon 1, Lyon, France; Department of Microbiology and Immunology, Penn State College of Medicine, Hershey, PA

**Keywords:** Hepatitis B virus, cccDNA, HBV replication, persistent HBV infection, cccDNA replenishment, de novo infection, anti-HBs antibody, chronic hepatitis B, HBV functional cure

## Abstract

Hepatitis B virus (HBV) infection is largely noncytopathic and requires the establishment of covalently closed circular DNA (cccDNA), which is considered stable in the nuclei of infected cells. Although challenging, approaches to directly target cccDNA molecules or kill infected cells are recommended to eliminate cccDNA. Herein, cccDNA levels were investigated in HBV-infected chimeric mice with humanized livers. HBV-infected cells support robust replication, progressively retain viral products, and head for cytopathic destruction and cccDNA loss. It is difficult for infected cells to retain cccDNA and remain noncytopathic. Replication-driven cccDNA loss is observed at both phases of spread of and persistent infection. The cccDNA replenishment is required to compensate for cccDNA loss. Blocking cccDNA replenishment pathways reduces cccDNA levels by >100-fold. These results prove an unconventional cccDNA elimination strategy that does not directly target cccDNA but aims to transform spontaneous cccDNA loss into progressive cccDNA elimination by blocking cccDNA replenishment.

## INTRODUCTION

Hepatitis B virus (HBV) is a hepatotropic DNA virus that may cause persistent and largely noncytopathic infection if caught in infancy^1,2^. The establishment and maintenance of HBV infection in hepatocytes require the formation of episomal covalently closed circular DNA (cccDNA) molecules, which function as templates for viral transcription in the nucleus^3,4^. Therefore, a single copy of cccDNA in an infected cell is minimally required. The cccDNA molecules are assumed to be long-lived^5^ because a chronic HBV infection usually lasts for years or decades^6^. Chronic HBV infection is thought to result from the failure of the host’s immune system to clear the established infection^7^; therefore, it is conventionally viewed as a continuation of the established initial infection. Current HBV cure strategies aim to directly eliminate or permanently silence cccDNA^5^, or clear infected cells from the livers^8^.

HBV replicates robustly in infected human hepatocytes, as evidenced by high serum hepatitis surface B antigen (HBsAg), HBV DNA levels^9–11^, and accumulated HBsAg and hepatitis B core antigen (HBcAg) proteins in infected cells during chronic HBV infection^12,13^ or in vitro infection^14,15^. The intracellular accumulation of viral products may indicate that the secretion of viral particles by infected cells lags behind HBV replication capacity, which may cause cytopathic changes if not stopped. The retention of L protein in hepatocytes of HBV transgenic mice causes a spectrum of pathologies, including necrosis and persistently elevated ALT levels, and the severity of pathology is related to the concentration of intracellular envelope proteins^16^. The intracellular HBsAg accumulation within smooth endoplasmic reticulum (ER) causes ER hyperplasia and displaces other organelles to the cell periphery, giving an appearance of “ground-glass” in some hepatocytes in chronic infection^17,18^. However, most HBsAg-positive cells in liver sections do not show a ground-glass appearance^19^. In the early 1990s, Summers et al. discussed the following principles on the replication and persistent infection of Hepadnaviruses: i. a persistent infection of Hepadnaviruses is contingent upon the inhibition of replication at late phase of infection; ii. control of cccDNA copy numbers is required to maintain persistent noncytopathic infection; and iii. intracellularly accumulated L protein acts as an overall suppressor of replication and permits persistent infection^20,21^.

Clinical evidence has shown the dynamic evolution of the cccDNA population. The wild-type viral population in serum or cccDNA in the liver can be cleared and replaced with mutant populations during chronic hepatitis B infection^22–26^. Complete cccDNA turnover in patients with chronic hepatitis B receiving nucleotide analog (NA) treatment occurs in a duration as short as 24 weeks^26^.

The loss of pre-existing cccDNA is supported by the quantitative detection of cccDNA levels in serial liver tissues, which showed that cccDNA levels were progressively reduced by 20-to 100-fold during NA treatment of woodchucks chronically infected with woodchuck hepatitis virus^27^, an animal model closely resembling chronic HBV infection in humans. A 1–2.9 log reduction in cccDNA levels has also been reported in human patients treated with NA^28–33^. The detected cccDNA reduction raises the possibility that cccDNA molecules can be spontaneously cleared from infected cells.

Taken together, we hypothesized that in vivo HBV-infected cells spontaneously clear cccDNA and tested this hypothesis in uPA/SCID chimeric mice with humanized livers that support a robust persistent HBV infection in the absence of functional T-and B-cell immunity^34^. Here, we present evidence supporting the concept of spontaneous cccDNA loss in HBV-infected cells and an unconventional cccDNA elimination strategy that aims to transform spontaneous cccDNA clearance into progressive cccDNA elimination by blocking cccDNA replenishment. The results herein identify cccDNA replenishment as the main cause for cccDNA persistence in HBV infection and provide proof-of-concept for an efficient cccDNA elimination therapy.

## RESULTS

Our strategy to test the hypothesis included three components:

1. Assessing the impact of efficient HBV replication on the presence of cccDNA in infected cells
2. Analyzing cccDNA levels at the bulk-cell and single-nucleus level
3. Evaluating therapeutic impact on cccDNA levels by blocking cccDNA replenishment

### In vivo replication kinetics suggests that the inhibition of HBV replication is likely through cccDNA clearance

To understand the kinetics of in vivo HBV replication, HBV-infected, untreated chimeric mice were euthanized on days 18, 45, 50, 52, 82, 99, 141, and 212 post inoculation (pi) for assaying kinetic serum HBsAg and HBV DNA levels and intrahepatic HBV markers. There were two infection phases (Figure 1A and 1B). The first was the phase of spread of infection to all infectible cells at which both serum HBsAg and HBV DNA levels rapidly increased after inoculation and peaked at around day 82 pi. The second was the persistent infection phase where HBV infection was maintained at a steady level. The observed kinetics of serum HBsAg and HBV DNA in this model recapture the typical acute HBV infection that becomes persistent in humans^9^.

**Figure 1.**
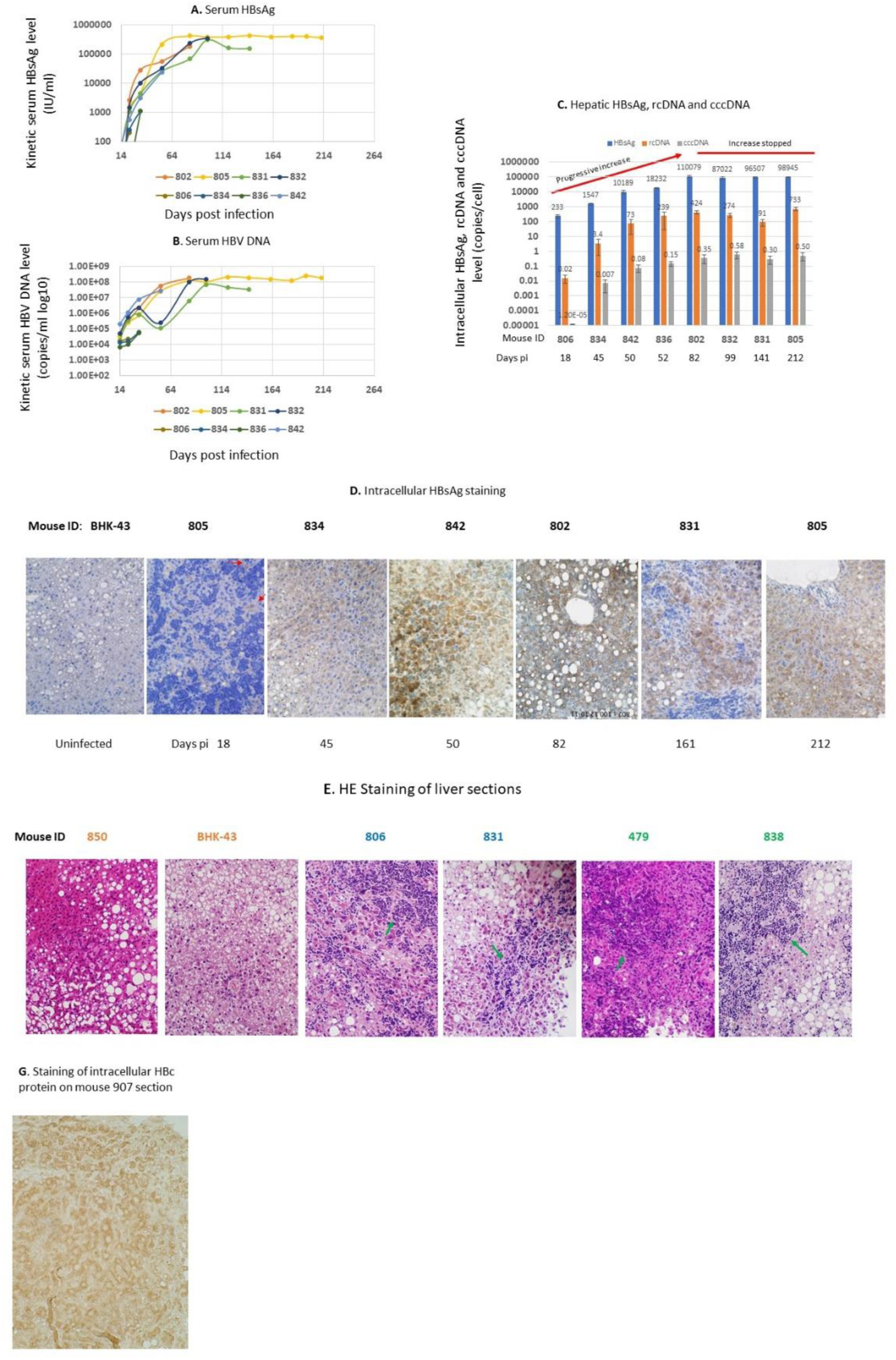
Intracellular accumulation of viral products is prevented from further increasing upon reaching the peak infection. **A.** and **B.** Kinetic serum HBsAg and HBV DNA levels. **C.** Intrahepatic HBsAg, rcDNA and cccDNA levels (copies/cell) were progressively increased during the spread of infection, then maintained at steady levels during persistent infection phase. Intracellular HBsAg was measured with ELISA. One mouse was sacrificed at each timepoint. **D**. Immunostaining of intracellular HBsAg shows the spread of infection by increasing the number of HBsAg positive cells and it appears that most cells were HBsAg positive around day 50 pi. Red arrows indicate two HBsAg positive cells on day 18 section. **E**. H&E staining shows severe liver parenchymal necrosis with infiltrates (green arrows) among HBV infected mice (orange: uninfected;: blue: untreated; green: treated). pi: post infection; cccDNA, covalently closed circular DNA; rcDNA, relaxed circular DNA. H&E, hematoxylin and eosin; HBsAg, hepatitis B surface antigen. Error bars were plotted with standard deviations.

The kinetics of intracellular accumulation of viral products also comprises two phases (Figure 1C). The first was the phase of a progressive increase in the accumulation of viral products. For instance, intrahepatic HBsAg levels increased from 230 copies per cell on day 18 to 110,000 copies on day 82 pi, which reflects a robust HBV replication and that the secretion of virions and subviral particles lags behind unrestricted HBV replication during the accumulation phase. The second was the phase of stopping the increase in the accumulation after the peak on day 82 pi. For instance, intracellular HBsAg levels remained around 100,000 copies/cell after the peak. Average cccDNA levels fluctuated 2-fold but also stopped increasing over the next 130 days. Further increases in HBV replication levels were likely suppressed during this phase because intracellular levels of viral products stopped increasing; however, serum HBsAg and HBV DNA levels remained steady (Figure 1A and B), indicating the stopped increase in the accumulation was unlikely caused by an increase in the secretion of viral particles.

A continuous accumulation of viral products can destroy the infected cells, causing cccDNA loss, which suggests that cccDNA cannot persist for long in the infected cells under such circumstances. Direct cytopathic effects of HBV infection in this model were reported^35^. Severely confluent liver necrosis with numerous infiltrates that involved up to 50% of the parenchyma on sections was observed in 4 of 18 HBV-infected livers (Figure 1E) but was absent in the remaining 14 liver sections (Figure S1). This suggests that the accumulated viral products trigger cellular responses to suppress cccDNA-driven HBV replication and avoid severe injury in most cells.

Our results suggest that in vivo cccDNA kinetics inckudes two phases (Figure 1C), and cccDNA may have been lost in both phases:

i. **Amplification phase**. The total cccDNA level in the liver was amplified mainly by expanding the infection in the liver. The cccDNA pool in individual infected cells is amplified through the intracellular recycling pathway in the early phase of infection^4,14,36,37^. The cccDNA level was increased from 0.00001 copies/cell (day 18 pi) to 0.35 copies/cell on day 82 pi. This was not the only event that occurred. Reaching peak infection means that all infectible cells must have been infected, as evidenced by the detection of HBsAg (Figure 1D) and HBcAg (Figure 2E) in almost all hepatocytes. The cccDNA level is expected to be ≥1 copy/cell because a minimum of one copy of cccDNA is required in each infected cell. However, the average cccDNA level at peak infection (day 82 pi) was 0.35 copies/cell, that is, approximately one copy of cccDNA per three infected cells, suggesting that cccDNA after the initial establishment may have been lost in a portion of infected cells.
ii. **Maintenance phase**. cccDNA was maintained at a steady level (0.35–0.6 copies/cell) to maintain HBV infection at a steady level upon reaching the peak. With an average cccDNA level of <1 copy/cell, the Poisson distribution predicted that some cells may contain >1 copy/cell and other cells may contain no cccDNA. This suggests that cccDNA was spontaneously cleared from some cells during the maintenance phase. Therefore, a steady HBV infection level in the persistent infection phase is likely reached by establishing an equilibrium between the number of infected cells with cccDNA that maintain HBV replication and those that have lost cccDNA and ceased viral replication. This implies that the infected cells regulate HBV replication mainly by clearing cccDNA.

**Figure 2.**
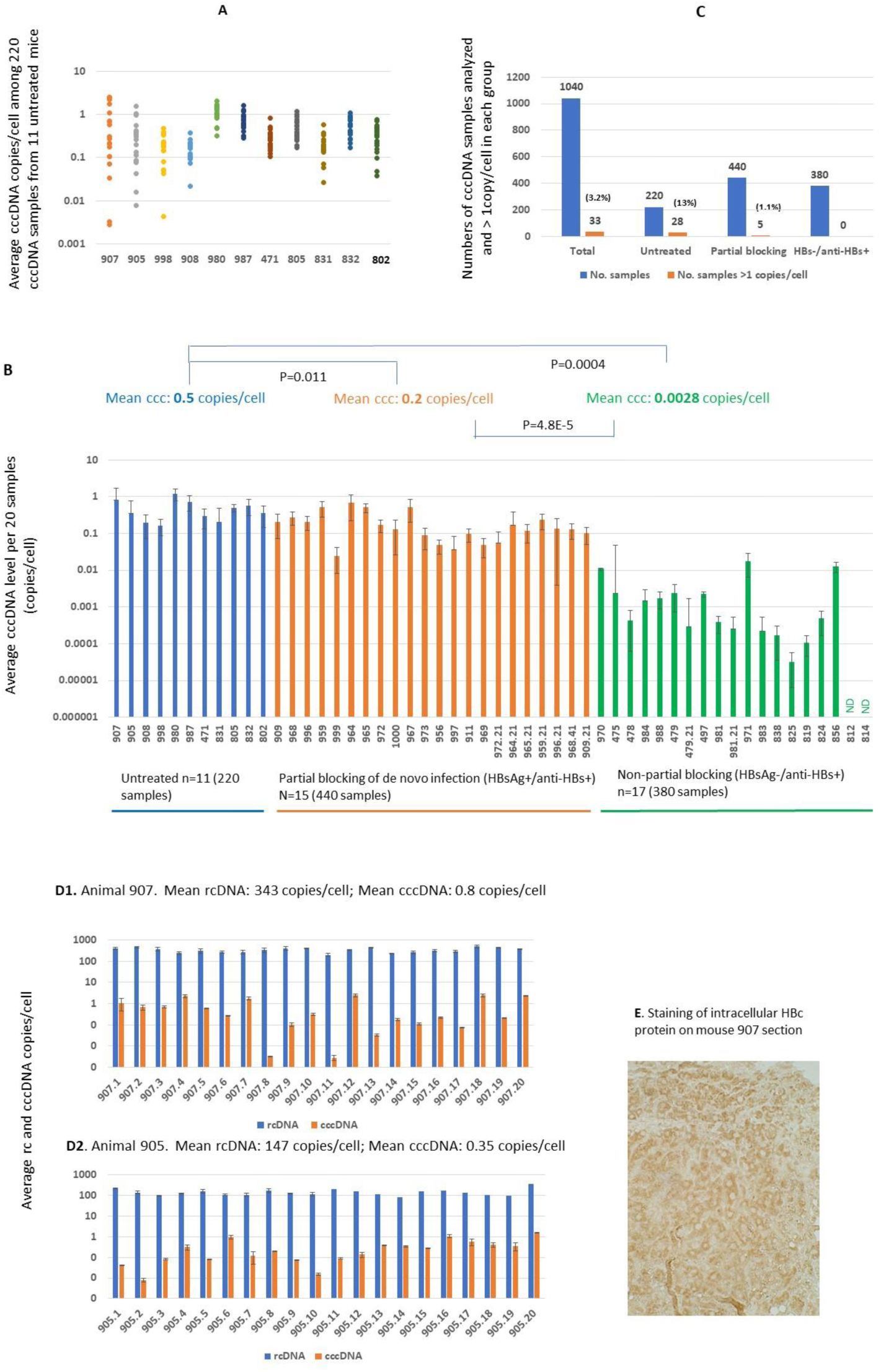
cccDNA levels in 192 of 220 cccDNA samples from 11 untreated mice were <1 copy/cell. **A.** Average cccDNA levels among 220 cccDNA samples. **B**. Average cccDNA level among 11 untreated livers (Blue) was 0.5 copies/cell. **C.** cccDNA levels > 1 copy/cell was detected in only 13% of 220 cccDNA samples. **D1** and **D2.** Diverse cccDNA levels (orange) among 20 cccDNA samples from the same livers despite comparable rcDNA levels (blue). **F**. Most cells were HBcAg positive (mainly in cytoplasm) on section of mouse 907 by immunohistochemical staining. ND: Not detectable; HBcAg, hepatitis B core antigen. The difference was considered significant at P<0.05 by Student’s *t* test. Error bars were plotted with standard deviations.

### Average cccDNA levels after peak infection were <1 copies/cell

We extended the analysis of cccDNA levels in untreated mice to obtain the range of cccDNA levels in livers. To avoid non-representative findings by a single sampling or a few samplings, we routinely sampled each liver 20 times, resulting in 220 cccDNA samples from 11 livers collected between days 82 and 253 pi. The highest average cccDNA level was 2.5 copies/cell while the lowest was 0.003 copies/cell among the 220 cccDNA samples (Figure 2A).

The average cccDNA levels in 28 (12.7%) of the 220 cccDNA samples were >1 copies/cell, whereas there were <1 copies/cell in the remaining 192 (87.3%) cccDNA samples (Figure 2B), indicating that some cells may not contain cccDNA molecules at different time points.

The detected cccDNA level per liver varied considerably from 1.2 to 0.16 copies/cell among 11 livers (Figure 2C). An average cccDNA level > 1 copies/cell (1.2 copies/cell) was detected in only 1 of the 11 livers.

The average cccDNA levels among the 20 samples from the same liver substantially differed. In animal 907 (Figure 2D1), the average cccDNA level in sample 12 and 8 was 2.5 and 0.003 copies/cell (one copy per 333 cells), respectively, with a >100-fold difference. However, rcDNA levels in two samples were comparable (343 vs 345 copies/cell). HBcAg staining showed that most cells were core protein-positive in the mouse 907 liver section (Figure 2E), suggesting that the low cccDNA level in sample 8 likely resulted from a scenario where cccDNA was already cleared to stop HBV replication but rcDNA remained in the infected cells because it took longer for more copies of rcDNA to be cleared after cccDNA clearance. In animal 905 (Figure 2D2), the cccDNA level in sample 20 was 1.53 copies/cell, while 0.01 copies/cell was observed in sample 2. The difference was also >100-fold though the difference in rcDNA level was <3-fold (364 vs 138 copies/cell). Different cccDNA levels among different samples from the same liver suggest a dynamic status of cccDNA in infected cells, that is, cccDNA molecules are maintained in some cells, but are lost in other cells in the same liver.

### cccDNA loss detected at a single nucleus level

One of the criteria used by the vendor PhoenixBio for selecting uPA/SCID chimeric mice with human livers is a liver replacement index (RI) of >70%^34^. Among the 57 mice received, only four had RI between 76–78% and ranged between 80–93% in the remaining 53 mice, approximately 76–93% of liver cells being human liver cells. As bulk cells are routinely used for cccDNA quantification in our assay, we used an average of 30% non-human liver cells to normalize the calculated cccDNA copies/cell. The number of non-human liver cells varied in each sample, which may have impacted the calculated copies/cell. We sought to quantitatively detect cccDNA copies at the single nucleus level^38^ to corroborate the absence of cccDNA in some infected cells. HBV rcDNA is 100–1000-fold more abundant than cccDNA (Figure 2D) and is also expected to be delivered to the nucleus for cccDNA conversion^37^. We simultaneously detected cccDNA and rcDNA in each nucleus using Absolute Q duplexing digital PCR (ABQ-duplexing dPCR), and the detected rcDNA was used as an HBV infection marker. The strategy, principle, and controls of the simultaneous detection of cccDNA and rcDNA are described in detail in the Methods section and Figure S2.

Nuclei from three livers, harvested on day 141, 218 or 253pi from untreated mice 831, 987, and 907, respectively (Figure 2A), were deposited in one nucleus per well in a 96-well plate using a BD FACSAria II. The total number of analyses performed is listed in Table 1.

**Table 1.**
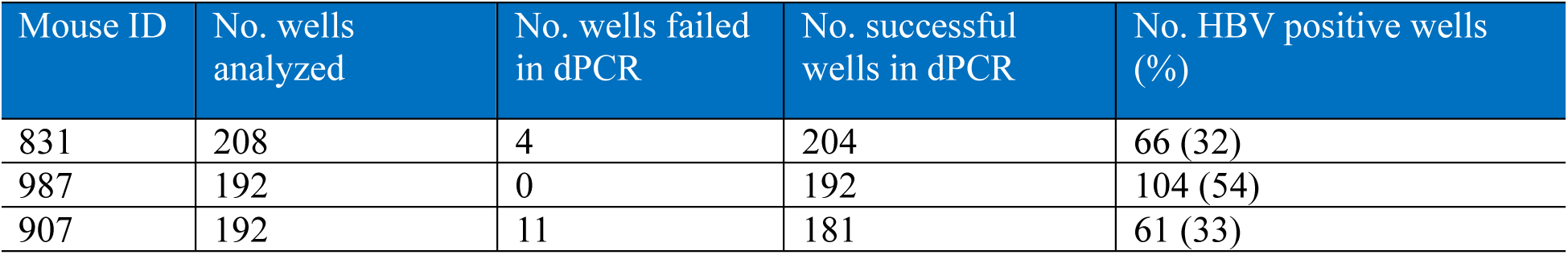
Percentages of HBV positive nuclei determined by duplexing dPCR.

| Mouse ID | No. wells analyzed | No. wells failed in dPCR | No. successful wells in dPCR | No. HBV positive wells (%) |
| --- | --- | --- | --- | --- |
| 831 | 208 | 4 | 204 | 66 (32) |
| 987 | 192 | 0 | 192 | 104 (54) |
| 907 | 192 | 11 | 181 | 61 (33) |

#### Detected cccDNA and rcDNA at the single nucleus level

cccDNA was detected as cccDNA only or coexisted with rcDNA (Figure 3A and B), and rcDNA was detected coexisting with cccDNA or rcDNA only (Figure 3B, E1B, E2B, E3B, and E4B).

**Figure 3.**
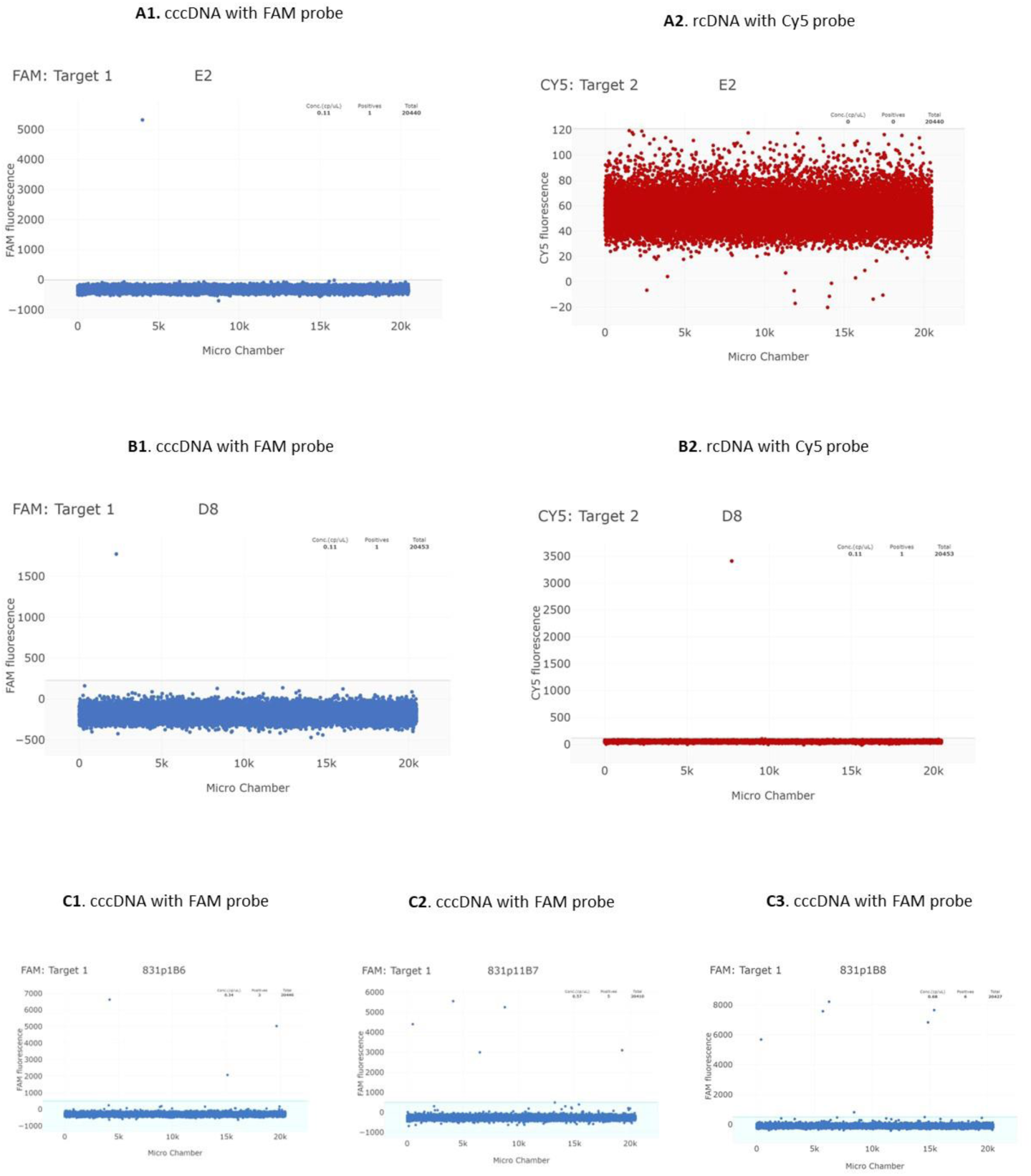
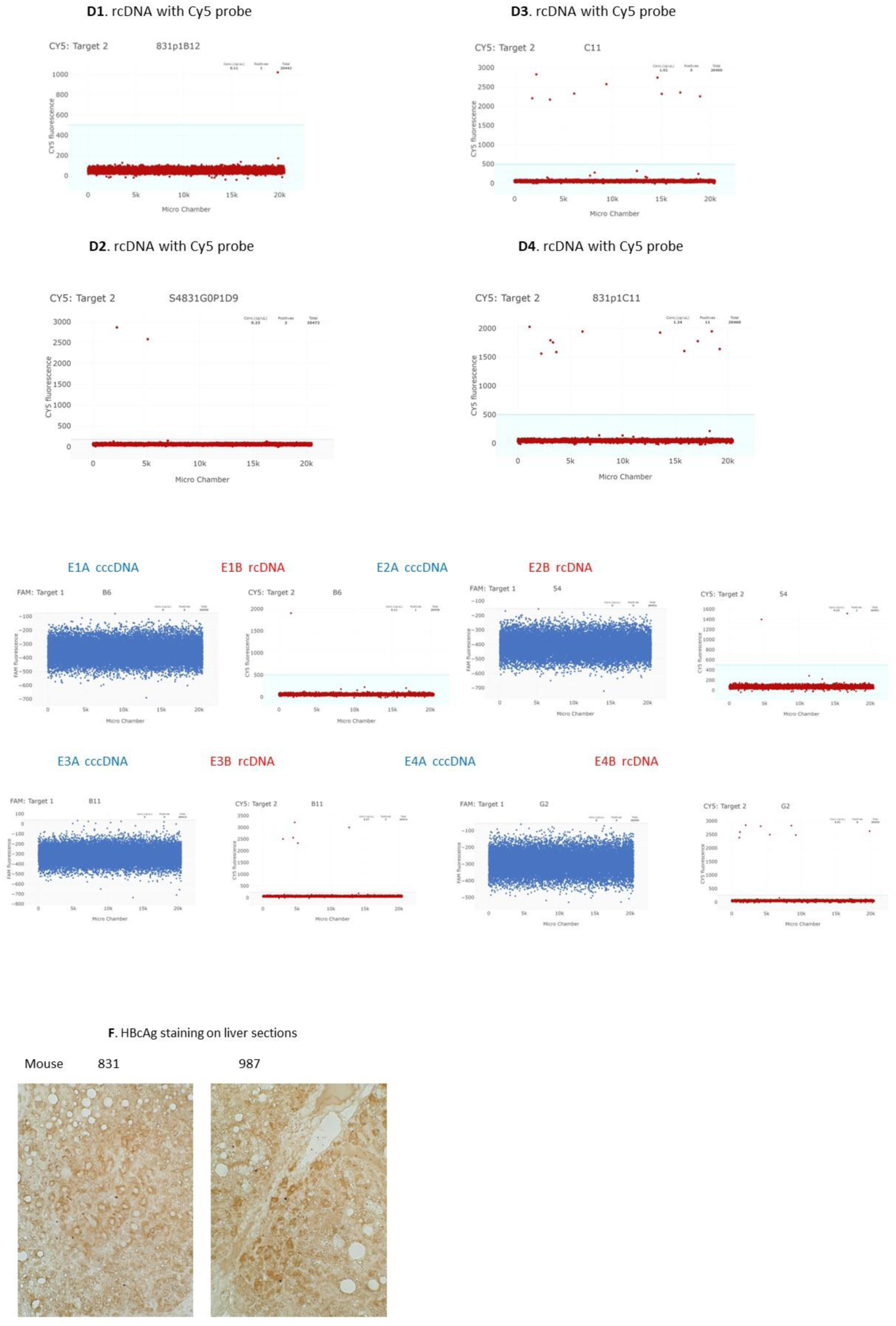
cccDNA and rcDNA detected at individual nucleus by ABQ duplexing dPCR. **A.** A single copy of cccDNA detected (A1) but no rcDNA detected (A2) in the same nucleus. **B**. A single copy of cccDNA (A2) as well as a single copy of rcDNA (B2) detected in the same nucleus. **C**. 3 (C1), 5 (C2), and 6 (C3) copies of cccDNA detected in 3 different nuclei, respectively. **D**. 1 (D1), 2 (D2), 9 (D3), and 11 (D4) copies of rcDNA detected in 4 different nuclei, respectively. **E**. cccDNA-/rcDNA+ nuclei. No cccDNA detected in E1A, E2A, E3A, and E4A while rcDNA detected as 1, 2, 5, and 8 copies in corresponding nuclei (E1B, E2B, E3B, and E4B). **F**. Immunohistochemical staining of HBcAg shows most cells were core protein positive on sections of mice 831 and 987.

#### cccDNA copies per nucleus

The cccDNA was detected as a single copy in most of the cccDNA-positive nuclei. Twenty (66.7%) of the 30 cccDNA-positive nuclei in mouse 831 contained only a single copy, while the remaining 10 nuclei had >1 copy, ranging from two to eight (Figure 3C). In mouse 987, 41 (75%) of the 55 cccDNA-positive nuclei contained a single copy of cccDNA, while 14 (25%) nuclei had > 1 copy. In mouse 907, the cccDNA was detected as a single copy in 34 (77%) of the 44 cccDNA-positive nuclei, while the remaining 10 nuclei contained >1 copy ranged between 2– 6 copies/nucleus. Thus, ≥2/3 of the detected cccDNA positive nuclei only contained a single copy of cccDNA, which is predisposed to lose.

#### rcDNA copies per nucleus

A single copy of rcDNA was detected in 24 (57%) of the 42 rcDNA-positive nuclei in mouse 831, and the remaining 18 (43%) nuclei had 2–11 copies/nucleus (Figure 3D). In mouse 987, a single copy of rcDNA was detected in 41 (60%) of the 66 rcDNA-positive nuclei, while the remaining 25 rcDNA-positive nuclei contained 2–8 copies/cell. In mouse 907, rcDNA was detected as a single copy in 13 (52%) of the 25 rcDNA-positive nuclei, and the remaining 12 (48%) nuclei had >1 copy of rcDNA, ranging from 2 to 19 copies/nucleus.

#### cccDNA-/rcDNA+ nuclei (Figure 3F)

A portion of infected cells from 27%, 47%, and 55% of the nuclei among the three livers (Table 2) did not have any detectable cccDNA, whereas rcDNA was detectable among the same nuclei (Figure 3E). Our infection kinetics data (Figure 1A and B) and the published data on HBV infection kinetics in this model^39^ show that peak infection can be reached on days 82–90 pi, implying that all infectible human liver cells are likely already infected at approximately 90 days pi. HBsAg and HBcAg staining showed that most cells were positive (Figure 2E, 3F, and 6D) in mice 831, 987, and 907 sections. Thus, cccDNA-/rcDNA+ cells likely represent either a loss of cccDNA from infected cells or cccDNA is yet to be formed with the delivered rcDNA from de novo infection in recently generated uninfected cells after cccDNA loss.

**Table 2.**
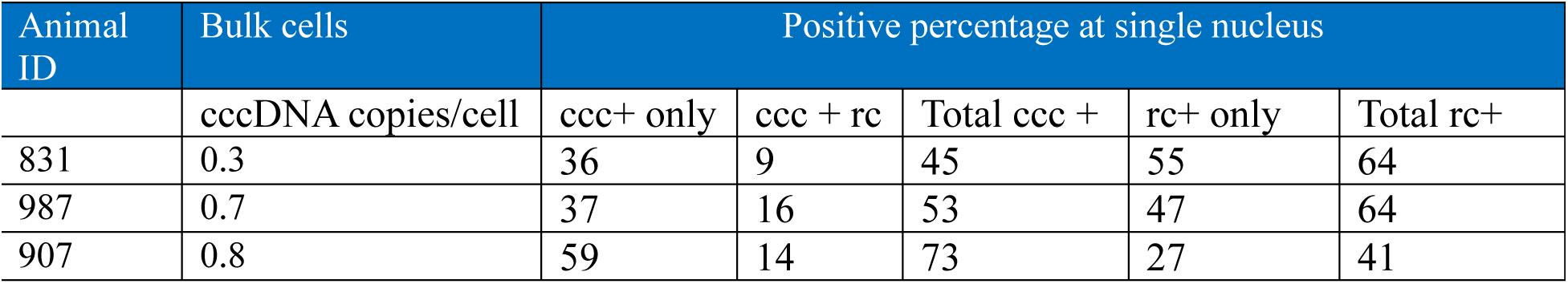
Percentages of cccDNA and rcDNA positive nuclei.

The detection of cccDNA-/rcDNA+ nuclei corroborated our bulk cell-based finding that cccDNA may have been spontaneously lost from a portion of the infected cells.

### Average cccDNA levels lowered by >100 fold upon blocking two cccDNA replenishment pathways

If the concept of spontaneous cccDNA clearance from infected cells is valid, then continuous cccDNA replenishment is required to maintain cccDNA levels. Therapeutic interventions aimed at blocking cccDNA replenishment should validate this hypothesis.

There are two known cccDNA replenishment pathways: intracellular recycling, which delivers newly synthesized rcDNA molecules into the nucleus for cccDNA conversion, and de novo infection^4,40^. Previous studies have suggested that de novo infection is the primary pathway of cccDNA replenishment^15,41,42^.

We first evaluated the therapeutic effect of blocking de novo infection with anti-HBs antibodies on cccDNA levels.

Two sources of anti-HBs antibody both of which are against “ad and ay” subtypes were used: exogenous mouse anti-HBs antibody (AM31509 PU-N OriGene) and endogenous anti-HBs antibody expressed by an AAV-anti-HBs vector HBVZ10 (described in detail in the Methods section). HBVZ10 can express high levels (up to 500 μg/mL) of anti-HBs antibody and was sustained at >100 μg/mL for at least 252 days in both immunocompetent and immunodeficient mice after a single injection (Figure S3A) or reached >100,000 mIU/mL if measured using WHO referenced calibrators, which are the standards for clinical report (Figure S3B).

A total of 15 mice received anti-HBs treatment, 13 of which were injected with the AAV-anti-HBs vector HBVZ10 at a dose of 1E11 genomic copies, and the remaining two with mouse anti-HBs antibody triweekly at a dose of 250 μg per injection (Figure S4A). Anti-HBs antibodies were detectable in all 15 chimeric mice after treatment. However, serum HBsAg remained positive (Figure S5A and S5B), suggesting that not all viral particles were neutralized, and de novo infection was only partially blocked.

The first set, consisting of six livers, and the second set, consisting of nine livers, were collected on days 204 and 253 pi, respectively. Each of the 15 livers was randomly sampled 20 times, and a second round of 20 samplings was performed in seven of the 15 livers, resulting in a total of 440 cccDNA samples. The average cccDNA levels in 440 samples are shown in Figure 4A. The average cccDNA levels per liver or per 20 samples are presented in Figure 2B (orange). The cccDNA levels >1 copies/cell were only detected in five (1.1%) of the 440 samples, which was significantly lower than the 13% of cccDNA samples in untreated mice. The mean cccDNA level in 15 mice with partially blocked de novo infection was 0.2 copies/cell, which is significantly lower (p=0.012) than 0.5 copies/cell in untreated mice (Figure 4B). Lower cccDNA levels in this group was significantly correlated with lower rcDNA levels (Figure 4C-4F). These results suggest that cccDNA levels are sensitive to partial blocking of de novo infection.

**Figure 4.**
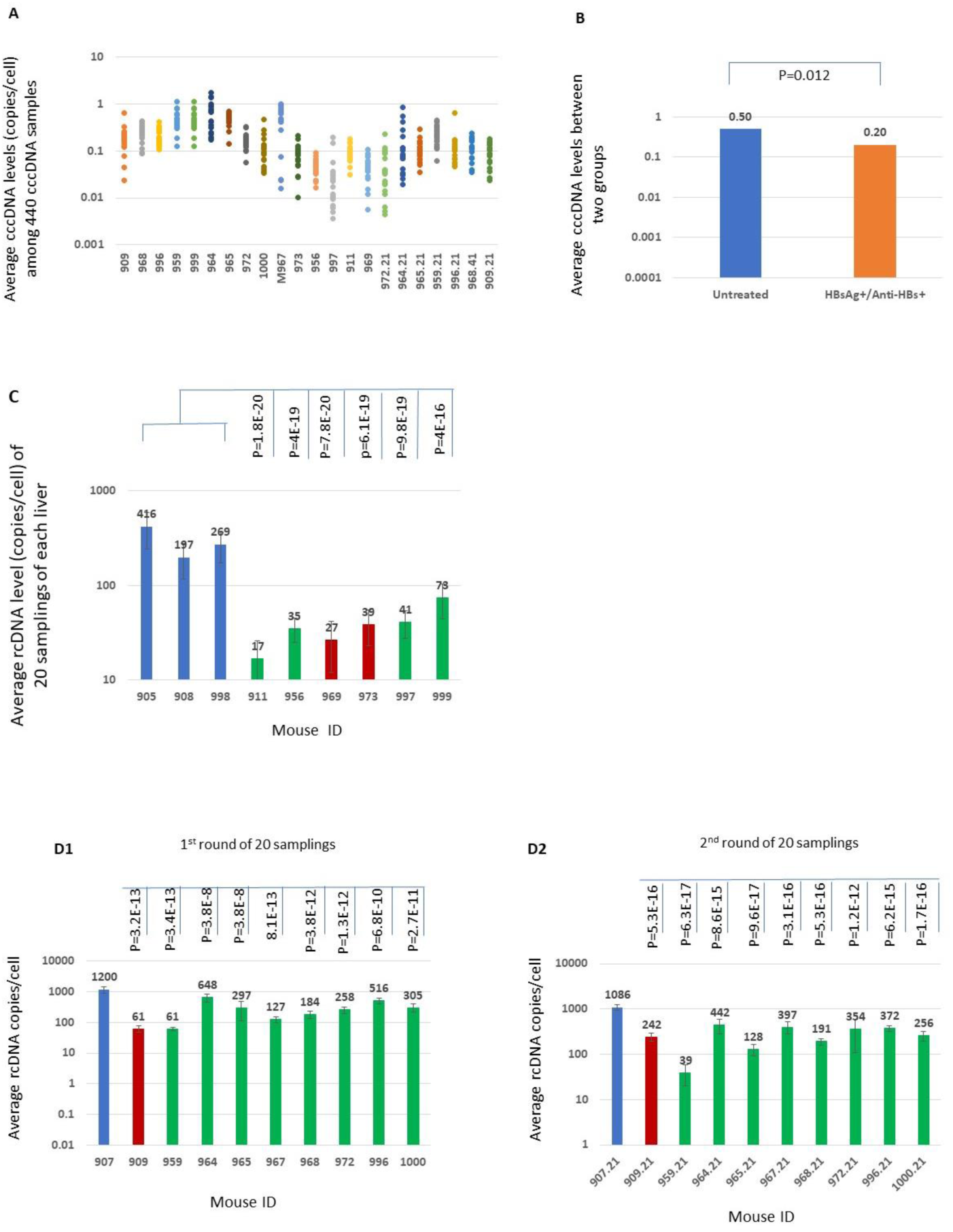
Partial blocking of de novo infection led to significantly lower cccDNA level. **A.** Average cccDNA levels were <1 copy/cell in 98.9% of 440 cccDNA samples isolated from the livers of 15 mice treated with the partial blocking of de novo infection. **B**. Average cccDNA level in this group was significantly lower by 60% compared to untreated mice. **C**. Average rcDNA levels significantly lowered in six livers treated with anti-HBs antibody harvested around day 204 pi compared to 3 untreated mice. **D**. Average rcDNA levels also significantly lowered among 9 mice treated with anti-HBs antibody compared to untreated mouse 907 (harvested on day 253 pi). The first and second round of 20 samplings for each liver (**D1** and **D2),** respectively. Blue: mock treated with AAV vector expressing malaria antibody. Green. Treated with AAV-anti-HBs vector HBVZ0 that expressed endogenous anti-HBs antibody. Red. Treated with exogenous mouse anti-HBs antibody. The difference was considered significant at P<0.05 by Student’s *t* test. Error bars were plotted with standard deviations.

We then evaluated the impact of non-partially blocking de novo infection or blocking both cccDNA replenishment pathways on cccDNA levels. The treatment schedule is shown in Figure S4B.

A non-partial blocking of de novo infection is marked by conversion of serum HBsAg from positive to HBsAg negative/anti-HBs positive (HBsAg-/anti-HBs+). Blocking of both cccDNA replenishment pathways was accomplished using a combination of anti-HB antibodies to block new rounds of infection with 9–12 weeks of entecavir therapy to decrease rcDNA synthesis and intracellular recycling of cccDNA (Figures S4B and C). A cccDNA analysis was performed on 17 mice that achieved HBsAg-/anti-HBs+ status (Figure 5C1, C2, D1, and D2), where the two mice (970 and 819) underwent anti-HBs antibody monotherapy, and the remaining 15 mice were treated with a combination of anti-HBs antibody and entecavir.

**Figure 5.**
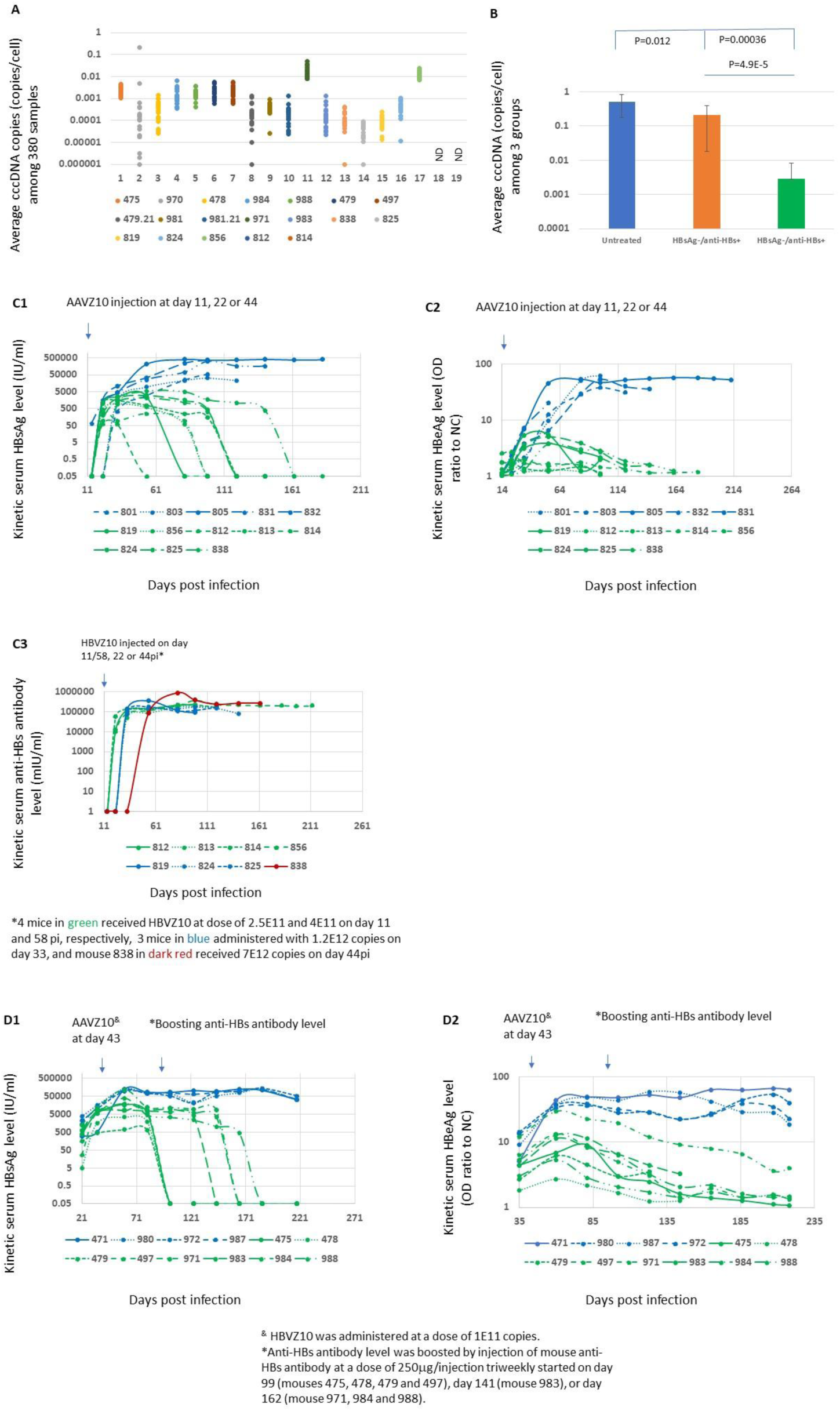
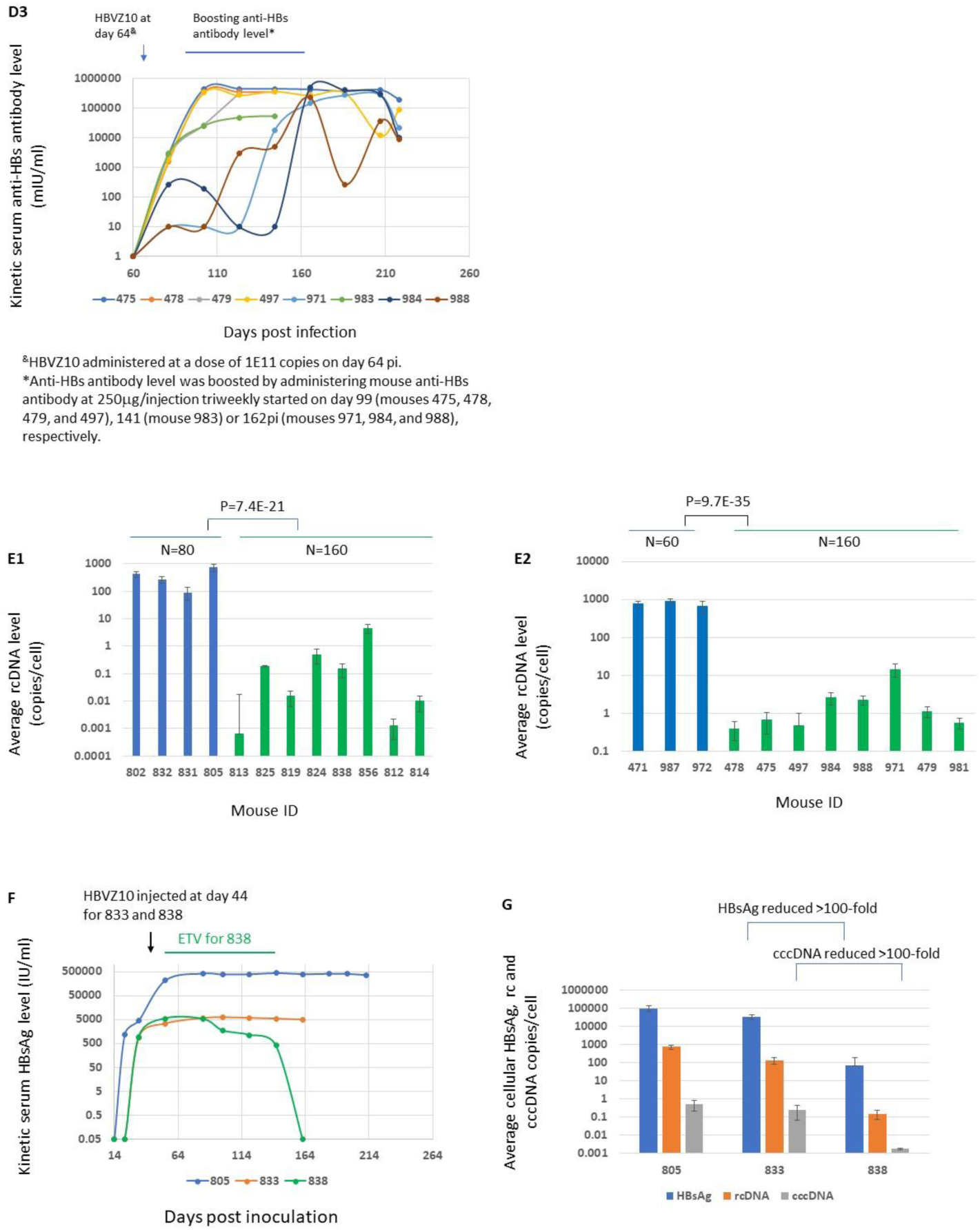
Non-partial blocking of de novo infection or blocking of two cccDNA replenishment pathways led to reducing cccDNA level by >100-fold in a few months. **A.** Average cccDNA levels were <1 copy/cell in all 380 cccDNA samples isolated from livers of 17 mice under non-partial blocking of de novo infection (two mice: 970 and 819) or blocking of two cccDNA replenishment pathways (the remaining 15 mice). **B.** Average cccDNA levels were significantly lower than both untreated and treated mice with partial blocking of de novo infection**. C1** and **C2.** Serum HBsAg and HBeAg levels were progressively reduced to undetectable levels among mice with non-partial blocking of de novo infection in mouse 819 or blocking of two pathways in the remaining 7 mice started before peak infection**. C3.** Kinetic serum anti-HBs antibody level**. D1** and **D2.** Serum HBsAg and HBeAg levels were progressively reduced to undetectable levels upon boosting anti-HBs antibody levels by triweekly administrations of mouse anti-HBs antibody at a dose of 250 μg/injection starting on day 99 pi or later until termination. **D3.** Kinetic serum anti-HBs antibody level in the same D1/D2 group. **E1** and **E2. Average rcDNA levels reduced by 2-4 logs among mice** through blocking of two pathways (Green) with anti-HBs antibody expressed by HBVZ10 before peak infection (E1) or with administering additional mouse anti-HBs antibody to boost the HBVZ10 expressed anti-HBs antibody level after the peak infection (E2) compared to untreated (Blue). **F**. Kinetic serum HBsAg levels among 3 mice (805 untreated; 833 HBVZ10 monotherapy, and 838 HBVZ10 combined with 12-week entecavir. **G**. Average intracellular HBsAg, rcDNA and cccDNA levels reduced by >100-fold in mouse 838 in 80 days compared to mouse 833. HBeAg, hepatitis B e antigen. The difference was considered significant at P<0.05 by Student’s *t* test. Error bars were plotted with standard deviations.

Each of the 17 livers that were collected after the peak infection between days 123 and 253 pi, were randomly sampled 20 times and a second round of 20 samplings was performed for the two livers, resulting in a total of 380 cccDNA samples (green in Figure 2B). Among these, 140 cccDNA samples from seven mice were analyzed using both qPCR and ABQ dPCR (Figure S6). All cccDNA levels were <1 copies/cell, and most were <0.01 copies/cell (Figure 5A). The mean cccDNA level in 17 mice was 0.0028 copies/cell, which is >100-fold lower (p=0.0001) than 0.5 copies/cell in untreated mice and significantly lower (p=4E-4) than 0.2 copies/cell in mice with partially blocked de novo infection (Figure 2B and 5B). In addition, cccDNA was not detected in two mice after 20 samplings of each liver. This demonstrates that non-partial blocking of de novo infection is critical for cccDNA elimination. The addition of entecavir to anti-HBs antibody blocks the recycling pathway and makes blocking of de novo infection by anti-HBs antibody more effective because the reduced virion production lowers the potential of de novo infection. These results further support the hypothesis that cccDNA replenishment is required to maintain cccDNA levels, underlining the spontaneous clearance of cccDNA from infected cells.

Kinetic human albumin levels were similar between untreated and treated mice (Figure S7), suggesting that cccDNA elimination mainly resulted from the blocking of cccDNA replenishment and not from the loss of human hepatocytes in humanized livers.

### cccDNA experienced progressive clearance upon blocking replenishment pathways in both infection phases

We further generated evidence supporting progressive cccDNA clearance upon blocking cccDNA replenishment pathways.

In HBV-infected chimeric mice, rising serum HBsAg levels paralleled rising viremia following inoculation (Figure 1A and 1B), suggesting that HBV replication is mainly cccDNA-driven and not integrated HBV DNA-driven. Entecavir treatment is known to reduce serum HBV DNA levels but does not have a direct impact on serum HBsAg levels, especially over a short duration^43^. Thus, serum HBsAg levels, but not HBV DNA levels, in entecavir-treated mice can be used as a surrogate for intrahepatic cccDNA levels. Serum HBsAg in all 16 mice (Figure 5C1 and D1) experienced a 3–5 log progressive reduction and became undetectable upon blocking the two cccDNA replenishment pathways. The blocking either started before peak infection, that is, during the cccDNA amplification phase (Figure 5C1) or the expressed anti-HBs antibody levels were boosted with the administration of additional doses of mouse anti-HBs antibody to turn partial blocking to non-partial blocking after the peak infection, that is, at the cccDNA maintenance phase (Figure 5D1). The progressive serum HBsAg reduction must have resulted from the progressive elimination of cccDNA. This was further supported by the progressive serum HBeAg reduction in 16 mice (green in Figure 5C3 and D3). HBeAg is synthesized using pre-core mRNA, which must be transcribed from cccDNA molecules^44^. The cccDNA level is generally reduced by 10–100-fold when HBV infection transitions from the HBeAg-positive to HBeAg-negative phase in chronic HBV infection^28,45^. Thus, a progressive serum HBeAg reduction (Figure 5C2 and 5D2) reflects the progressive reduction of cccDNA in the liver. Anti-HBs antibodies and entecavir do not directly target cccDNA molecules, therefore, the observed cccDNA elimination was likely mediated through spontaneous cccDNA clearance (non-treatment-mediated) that occurred in both phases.

### Efficient cccDNA clearance led to >100-fold reduction in cccDNA level in 80 days

uPA/SCID chimeric mice with human livers are fragile and cannot withstand stressful procedures, such as serial liver resections, posing a challenge in establishing baseline cccDNA levels before treatment. Therefore, cccDNA levels from different mice with comparable serum HBsAg levels were used as a reference. Figure 5F shows that baseline HBsAg levels on days 54 and 82 reached approximately 5000IU/mL before a progressive decline in mouse 838, which was treated with anti-HBs antibody expressed by HBVZ10 and entecavir. This is comparable to that in mouse 833 who received HBVZ10 monotherapy on day 44. Mouse 833 displayed dual positivity for serum HBsAg (Figure 5F) and anti-HBs antibodies (not shown) after treatment; thus, de novo infection in mouse 833 was considered partially blocked. Serum HBsAg levels remained steady at approximately 5000IU/mL from day 82 and 162 (termination day) in mouse 833. We used the cccDNA levels in mouse 833 as a reference for baseline cccDNA levels before HBsAg clearance in mouse 838. Both the intracellular HBsAg and cccDNA levels in mouse 838 were reduced by >100-fold during 80 days from day 82 to day 162 pi compared with those in mouse 833 (Figure 5G). The average reduction in cellular HBsAg levels was 423 copies/day. The cccDNA results further support that kinetic serum HBsAg levels can be used as a surrogate for cccDNA levels, and that the observed spontaneous cccDNA clearance was efficient and could be transformed into progressive cccDNA elimination upon blocking both cccDNA replenishment pathways.

### Confirmation of HBsAg reduction in both serum and liver with ELISA, western blot, and immunohistochemical staining

Two serial serum samples from untreated mice and two from treated mice, whose serum HBsAg underwent progressive reduction and became undetectable (Figure 6A1 and B1), were subjected to western blot analysis. This showed similar kinetics as detected by ELISA, confirming the progressive reduction in serum HBsAg in two serial serum samples with anti-HBs treatment (Figure 6A2, B2, and B3).

We also analyzed intrahepatic HBsAg levels in seven mice with progressive serum HBsAg reduction. Intrahepatic HBsAg was either not detectable or was detectable at low levels by ELISA (Figure 6C1), which was confirmed by western blot analysis (Figure 6C2).

Consistent with ELISA and western blot data, immunohistochemical staining of HBsAg in sections (Figure 6D) showed that intracellular HBsAg was reduced to undetectable or barely detectable levels among seven mice that achieved HBsAg-/anti-HBs positive status.

**Figure 6.**
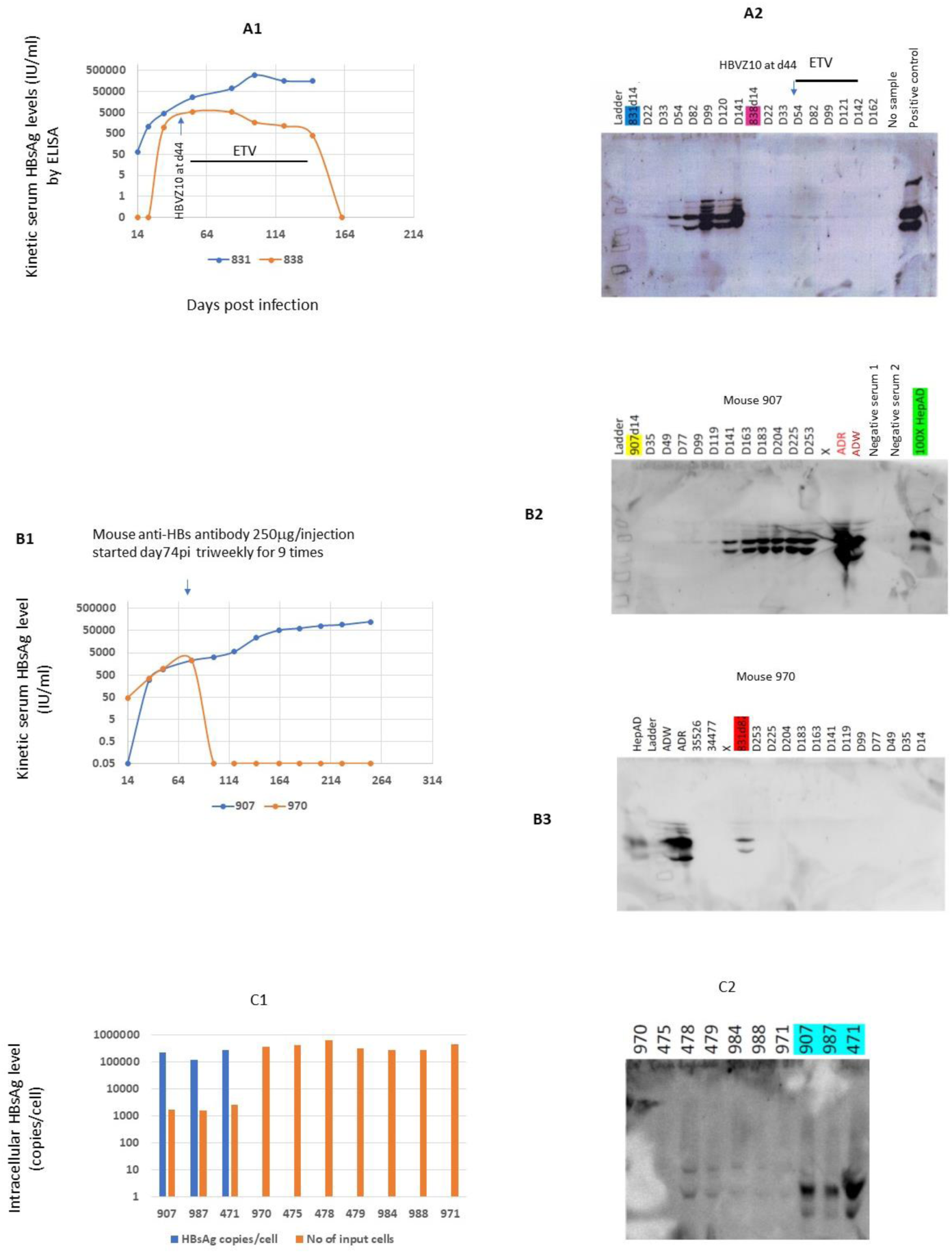
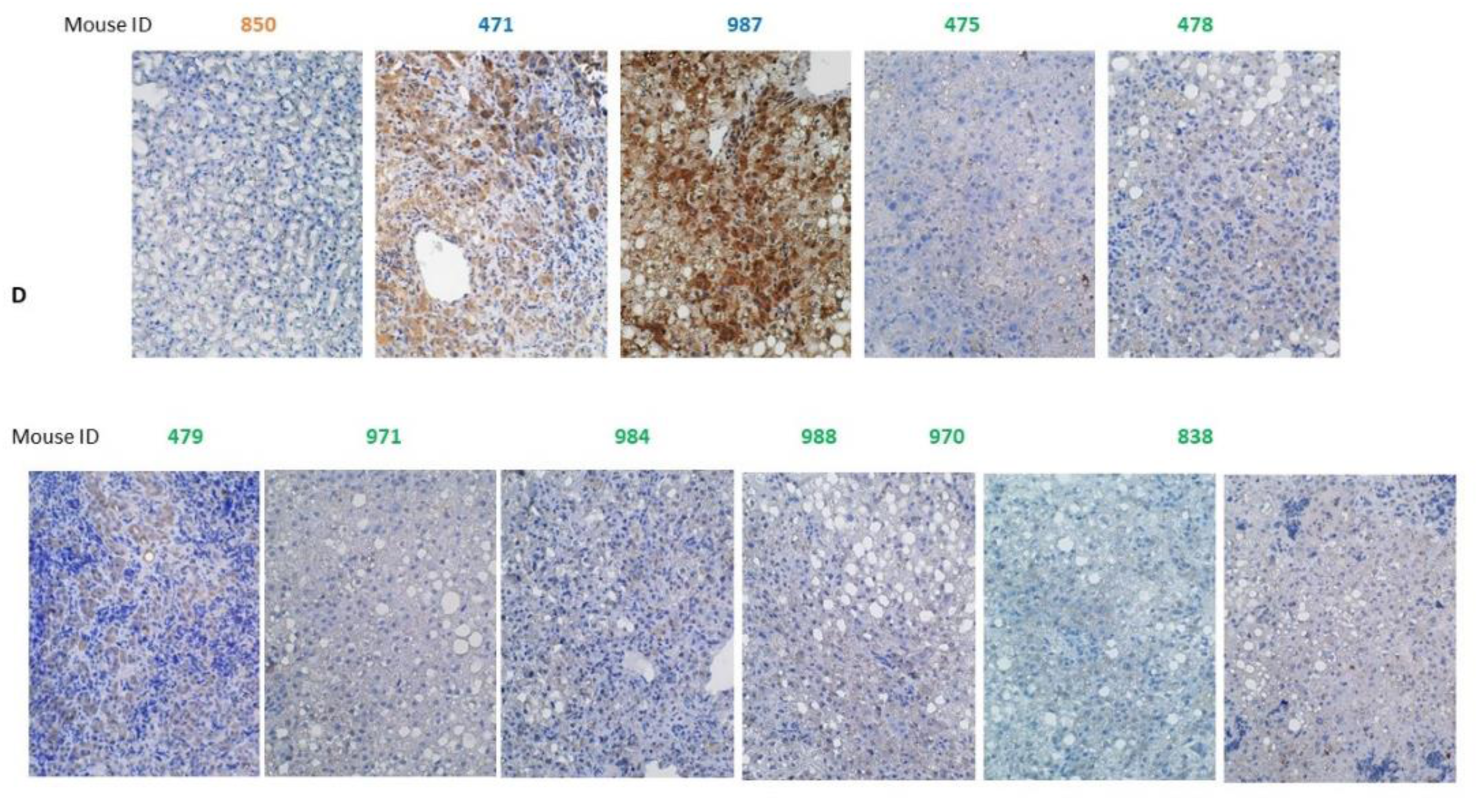
Progressive reduction in serum HBsAg detected by ELISA corroborated by western blots and supported by intracellular HBsAg reduction. **A.** Progressive reduction of serum HBsAg detected by ELISA in mouse 838 (A1 orange) who received a single dose of HBVZ10 combined with 12-week entecavir therapy and corroborated by western blot (A2). Mouse 831 was an untreated control. 100xHep: Positive control with concentrated viral particles from the HepAD38 cell medium. **B**. Progressive reduction of serum HBsAg detected by ELISA in mouse 970 (B1 orange) who received mouse anti-HBs antibody monotherapy at a dose of 250 μg/injection triweekly for 9 consecutive times and corroborated by western blot (B2). Mouse 907 was an untreated control. ADR and ADW: HBV positive human sera. 100xHepAD, Positive control with concentrated viral particles from HepAD38 cell medium. 35526 and 34477, negative human sera. X, no sample lane. Intrahepatic HBsAg was not detectable (C1) by ELISA that analyzed 30 μl of each lysate in 7 mice with HBsAg-/anti-HBs+ status and was either undetectable or significantly reduced in 7 livers compared to 3 untreated mice (mice 907, 987, and 471) by western blot (C2) that analyzed 40 μl of each lysate. **D**. Intracellular HBsAg reduced to barely detectable or undetectable among 8 mice (mouse ID in green) whose serum HBsAg was progressively reduced upon blocking of two cccDNA replenishment pathways, compared to 2 untreated mice (Blue in mouse ID). Mouse 850: uninfected.

## DISCUSSION

In contrast to the dominant view that HBV cccDNA molecules are stable in infected cells^5^, spontaneous cccDNA loss was observed during both the spread of infection and persistent HBV infection phases in this study using chimeric mice with humanized livers. Moreover, cccDNA replenishment was required to maintain cccDNA levels.

The analysis of cccDNA copies at the single nucleus level shows that most infected cells contain a single copy of cccDNA; nonetheless, average intracellular HBsAg levels are accumulated at approximately 100,000 copies/cell upon reaching the peak, highlighting an extraordinary efficiency of both RNA transcription and viral protein synthesis. The continuous presence of cccDNA after reaching the peak in infected cells is expected to continuously drive-up transcription and further increase the accumulation of viral products to an intolerable level, leading to cytopathic destruction through which cccDNA will be lost. Furthermore, unlike HIV, HBV is not known to have a latent infection phase, implying the absence of the noticeable inhibition of RNA transcription, which is supported by high serum HBsAg levels in both HBeAg positive and negative phases^10,11^. Thus, the long-term presence of cccDNA in infected cells may not be feasible under such circumstances.

However, HBV is largely noncytopathic, implying that infected cells may respond well to the stress caused by accumulating high levels of intracellular viral products. As reported during the second phase of hepadnaviral replication, viral-envelope proteins accumulate in the infected cells. The L protein alone or in combination with M and/or S proteins inhibits further cccDNA amplification or decreases cccDNA levels^46–48^. Because the highly efficient RNA transcription and viral-protein synthesis suggest the absence of efficient inhibition of these steps, clearing cccDNA appears to be the only effective option to stop HBV replication and protect cells from being destroyed.

Therefore, such replication-driven cccDNA loss can occur either through spontaneous clearance (Figure 7A) or cell destruction (Figure 7B). Cytopathic effects in infected primary hepatocytes and acute liver injury were also observed during in vivo infection with the duck hepatitis B virus (DHBV, a member of the Hepadnaviridae family) L protein mutant G133E, which caused a defect in enveloped virus production and increased intracellular levels of cccDNA, RNA, capsid, and rcDNA^20,49^.

**Figure 7.**
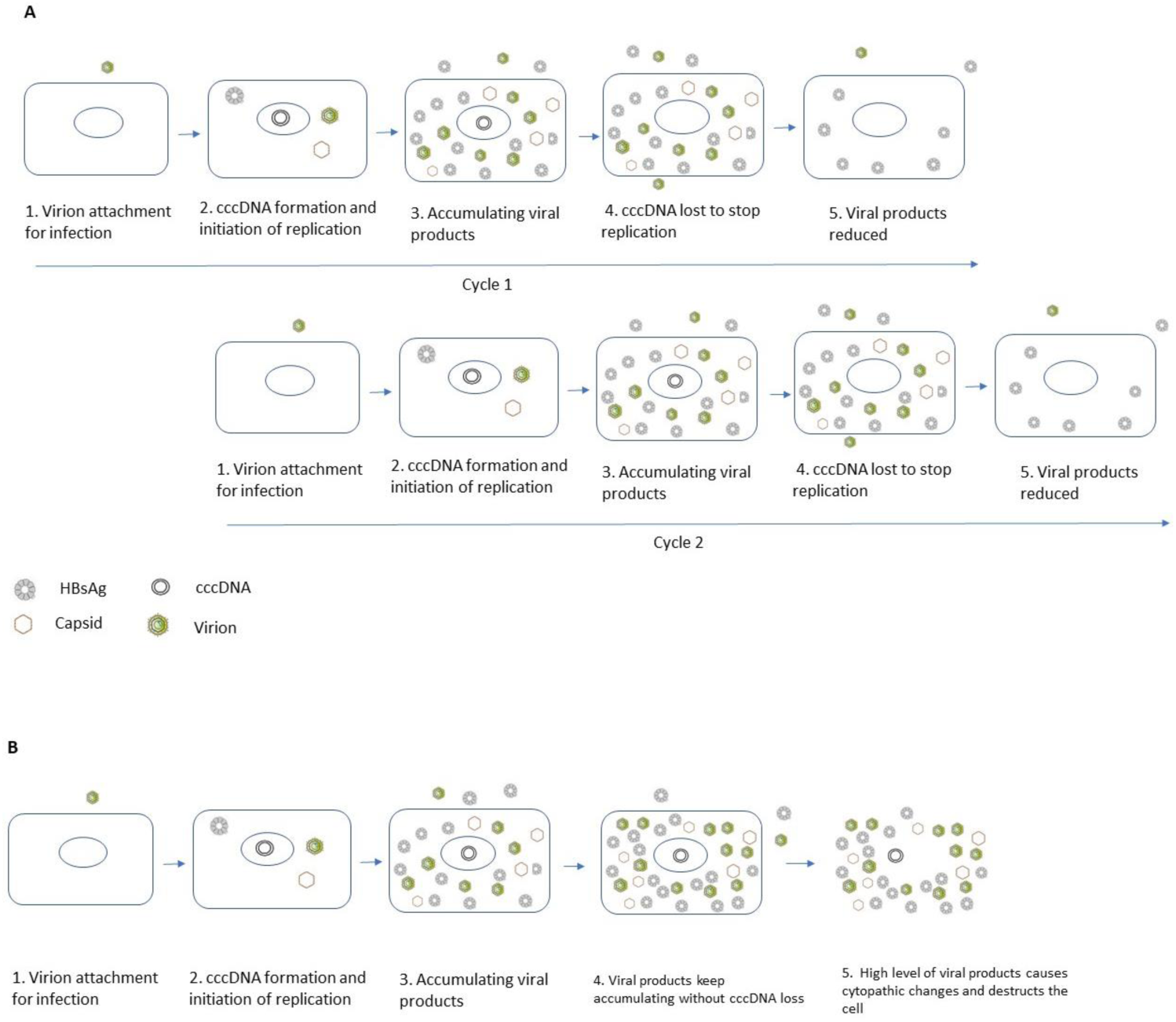
Proposed mechanisms for replication-driven cccDNA loss and persistent HBV infection. **A.** cccDNA loss triggered by the high level of accumulated viral products and the cyclic features of cccDNA dictates that persistent HBV infection consists of multi-cycle infection. **B**. cccDNA may eventually get lost through cell destruction if cccDNA is retained upon the accumulation of a high level of viral products.

The time course from cccDNA establishment and an initial amplification through the intracellular recycling pathway to cccDNA loss in infected cells represents one cycle of infection, suggesting that cccDNA in infected cells is only present for one cycle of infection: the cccDNA lifespan is approximately equivalent to the duration of a cycle of infection. A new cycle of infection is initiated when cccDNA is re-established through the de novo infection of HBV-negative cells. This cyclic feature of cccDNA indicates that persistent HBV infection is maintained by multiple cycles of de novo infection (Figure 7A).

Published studies on kinetic intracellular viral product levels showed that intracellular HBcAg reached a peak level at approximately 2–4-week in HBV infected HepG2-NTCP cells and was dependent upon the efficiency of viral particle secretion from infected cells^14,15^. In in vivo DHBV infection, the intracellular L protein reached a peak around day 5 pi^50^, suggesting that intracellular hepadnaviral infection may reach its peak at 1–4 weeks pi on average. In this timeframe, after which cccDNA clearance may occur, each cycle of HBV infection likely ends with cccDNA loss within a few weeks at the individual cell level. However, HBV infection at the liver level in the absence of sufficient anti-HBs antibodies may last for years or decades because there are always new cycles of infection, whereas early cycles of infection end following cccDNA loss.

Previous studies have suggested that cccDNA molecules may be lost during cell division^38,42^. Thus, spontaneous cccDNA loss could also be caused by random liver-cell turnover, which may occur at any stage. Because there is a clear pattern of progressive accumulation of intracellular viral products, cccDNA loss may mainly occur after reaching the peak infection in individual cells. Thus, we assume that the replication-driven cccDNA loss is more prominent than random cell turnover-mediated cccDNA loss.

Our findings prompt us to rethink the cccDNA elimination strategy. The direct targeting of cccDNA or killing infected cells is recommended for cccDNA elimination and complete cure of chronic HBV infection. Such a strategy may not be required based on this study. Instead, our results devise an unconventional cccDNA elimination strategy that does not require direct targeting of cccDNA molecules but aims to transform spontaneous cccDNA loss into progressive cccDNA elimination by blocking cccDNA replenishment.

A limitation is that this study did not investigate the contribution of human liver-cell proliferation to the detected cccDNA loss in this model. The proliferation rates of human liver cells in uPA/SCID humanized livers was reported to progressively reduce from the initial 68% on day 5 to 3% on day 100 post-human liver-cell transplantation^42^ when the growth of human liver was close to completion, suggesting that the proliferation rate of human liver cells is mainly determined by the available liver space for growth. Thus, the smaller the space, the lower the proliferation rate. The same previous study also observed that most HBcAg positive cells were non-proliferating cells, suggesting that non-proliferation is likely required to establish fully-fledged HBV replication in infected cells because cell division may cause cccDNA loss and abort replication at nascent stages. In this study, all infection experiments started approximately 120– 150 days after transplantation when the growth of the human liver was completed, and the proliferation of human liver cells was considered negligible (personal communication with PhoenixBio). The kinetics of in vivo replication observed in this model (Figure 1A-C) resemble a typical acute HBV infection that becomes chronic in humans^9^, suggesting that the proliferation of infected cells was not sufficiently frequent to interrupt HBV replication kinetics in this model. This observation is consistent with the findings that most cells on liver sections harvested after peak infection from untreated mice were HBsAg or HBcAg positive, which are likely non-proliferating cells^42^. Importantly, our results indicate that cccDNA can still be lost through the replication-driven mechanism from infected cells that supports the highly efficient replication and progressive accumulation of viral products despite the non-proliferation of the cells. Consequently, the findings herein will prompt the initiation of new studies on the relative contributions of replication-and cell division-driven mechanisms to cccDNA loss.

In summary, most HBV-infected cells in chimeric mice contained a single copy of cccDNA, but produced, accumulated, and maintained approximately 100,000 copies of surface protein per cell, reflecting the high efficiency of both RNA transcription and viral protein synthesis and the inability to secret all viral products of infected cells. Thus, cccDNA cannot be retained for long in the infected cells under such circumstances. We observed spontaneous cccDNA loss from infected cells, and cccDNA loss was compensated for by cccDNA replenishment. Our results devise an unconventional cccDNA elimination strategy that does not require direct targeting of cccDNA but aims to transform spontaneous cccDNA loss into progressive cccDNA elimination by blocking cccDNA replenishment pathways. Once translated in human trials, this cccDNA elimination strategy will deliver an efficient HBV cure treatment.

## Acknowledgements

We would like to thank Dr. David Baltimore at the California Institute of Technology for his generous support in developing this AAV vector-based HBV therapy; Dr. Rajen Koshy and Dr. Mindy Davis at National Institute of Allergy and Infectious Diseases (NIAID), National Institutes of Health (NIH) for participation, advice, and support of this project; Dr. Alex Balazs at Broad Institute for technical support; Ken Class at the University of Maryland for sorting nuclei; Dr. Todd Parsley at Noble Life Sciences; and Ms. Amanda Ulloa at NIAID for reading the manuscript. This study was supported by NIH Research Contracts 75N93020C00045 and 75N930220C00042, SBIR grant R43 AI155066-01A, and DoD Discovery Award W81XWH-21-1-0065 (YYZ). FZ received public grants overseen by the French National Research Agency (ANR) as part of the second “Investissements d’Avenir” program (reference: ANR-17-RHUS-0003) and by the European Union (grant EU H2020-847939-IP-cure-B). JH was funded by a NIH grant, R37AI043453.

## Author Contributions

Conceptualization: YYZ

Methodology: BHZ, FZ, JH and YYZ

Investigation: BHZ, YZ, SKH, HP, LL, JH and YYZ

Funding acquisition: YYZ

Project administration: SKH and BT

Writing – original draft: YYZ

Writing – review & editing: BT, FZ, JH and YYZ

## STAR METHODS

### Animals and HBV infection

All animal experiments were performed at Noble Life Sciences Inc (Sykesville, MD), a preclinical research contract service provider. The selection of Noble Life Sciences Inc as the subcontractor to perform animal experiments was approved by the NIAID contract office at NIH (NIH approved Animal Welfare Assurance number, A4633-01). All animal studies were approved by the Institutional Animal Care and Use Committee (IACUC) of Noble Life Sciences protocol NLS-614. All animals received humane care.

Immunocompetent female mice (CD1) were purchased from Charles River Laboratories (Boston, MA, USA), and immunodeficient male mice (uPA/SCID chimeric mice) were supplied by PhoenixBio USA (New York, NY, USA). All mice were kept in housing cages (TP107, One Corporation, Osaka, Japan) in an BSL-2 room with controlled temperature at 23°C and 12 hour-light/dark cycle. All animals fed with γ-radiated CRF1 food and autoclaved water *ad libitum*. An HBV inoculum, prepared from mouse serum (project no H01-108 animal 4) by diluting viremia of 5E9 HBV DNA copies/mL to 2E7 HBV DNA copies with PBS in 100 μl volume, was administered intravenously (tail vein) to each chimeric mouse.

In total, three HBV infected animal experiments were performed for this study; the time course and procedures of each experiment were illustrated in Figure S4.

### AAV-anti-HBs vector (HBVZ10)

HBVtech has developed a new HBV therapy candidate called AAV-anti-HBs vector, which utilizes an optimized adeno-associated virus (AAV) vector^51–54^ to deliver human anti-hepatitis B surface antigen (anti-HBs) antibody genes. Such AAV-anti-HBs vectors can express sustained high level of anti-HBs antibody after a single injection. The purpose of developing the AAV-anti-HBs vector as a new HBV therapy candidate is to remedy the deficiency in anti-HBs antibody production in chronic HBV infection. HBVZ10 is the leading AAV-anti-HBs vector and has been intramuscularly administered in chimeric mice to express human anti-HBs antibody endogenously and to block de novo infection in the absence or presence of entecavir.

### Production of AAV-anti-HBs or anti-malaria antibody vectors

Briefly, 293 cells were co-transfected with an AAV vector encoding anti-HBs or anti-malaria antibody and the plasmid pDP8.ape (PlasmidFactory, Bielefeld, Germany), which provides pHELP plasmid function and encodes AAV2 rep and AAV8 cap proteins in trans for packaging AAV vectors. The resultant AAV vector does not contain any viral open reading frame (ORF). AAV was purified via PEG precipitation and cesium chloride ultracentrifugation. The infectivity of AAV aliquots was confirmed *in vitro* by transducing 293 cells and quantifying the antibody concentration in the medium using ELISA. A total of 1E14 genome copies were obtained for each vector after production, purification, and concentration.

### HBVZ10 administration doses

A small dose of HBVZ10 at 1E11 genomic copies was intramuscularly administered in animal experiments 1 and 2 (Figure S4A and B), and the higher doses of HBVZ10 at 2.5E11, 1.8E12 or 7E12 genomic copies were administered for animal experiment 3 (Figure S4C).

### Monitoring HBV infection and human albumin level in blood

Blood was collected tri-weekly for quantification of serum HBV DNA (qPCR see below), HBeAg (CSB-E13557h, CUSABIO), HBsAg (GS HBsAg EIA 32591, Bio-Rad), anti-HBs antibody (MONOLISA Anti-HBs EIA 25200, Bio-Rad) with calibrators (MONOLISA Anti-HBs 20-Calibrator kit 25219, Bio-Rad) and human albumin (Human albumin ELISA kit E-80AL, Immunology Consultants Laboratory) levels by ELISA per instructions.

In addition, serum HBsAg in the selected samples were analyzed by western blot.

### Analysis of intrahepatic HBV DNA

Each liver was randomly sampled 20-40 times by cutting 20-40 mg liver tissue (weighed and recorded) and placed in a disposable micro-homogenizer (BioMasher, Takara cat no: 9790B) in 500 μl of an isotonic buffer (154 mM Tris-HCl, pH 7.5, 1 mM EDTA and 0.05% TritonX-100) with 10 strokes. The homogenized tissue suspension was spun for 2 min at 14,000 rpm and 100 μl lysate were saved for western blot of intracellular HBsAg and the remaining 400 μl transferred to a new microtube for isolation of replicative intermediates (RI) while nucleic pellet remained in the tube for cccDNA isolation.

Note: Two negative controls were included for each round extraction, one placed in the 1^st^ sample position and the other in the last position to monitor any contamination during extraction.

### Extraction of rcDNA from the 400 μl supernatant by following procedures^50^

1. Add 110 μl of proteinase K (final 0.5 mg/mL) with 1% SDS and incubate at 50°C for an hour.
2. Add 500 μl phenol, vortexed and chilled on ice for 3 min and centrifuge at 14,000 rpm for 2 min.
3. Transfer supernatant to a new tube and add 1000 μl of 100% ethanol for precipitation and centrifuge samples at 14,000 rpm for 15 min.
4. Wash pellets with 1000 μl of 100% ethanol at 14,000r pm for 10 min.
5. Remove residual ethanol and air dry for 5 min.
6. Dissolve pellet in 200 μl of 10:1 TE buffer, pH 7.4, and then the rcDNA is ready for qPCR

### Extraction of cccDNA from nucleic pellets by following procedures^38^

1. Suspend pellet with 200 μl of 10:1 TE with 0.05% Triton-X100 pH7.4.
2. Add 200 μl of 6% SDS-0.1 M NaOH solution and incubated at 37°C for 15 minutes.
3. Add 100 μl of 3 M KAc pH 5.07 and mix thoroughly. Chill on ice for 5 minutes and then microfuge to remove KSDS-protein-ssDNA complexes (pellet) at 14,000 rpm for 2 min.
4. Transfer supernatant to a new tube, add 500 μl of phenol, and centrifuge at 14,000 rpm for 2 min.
5. Recover supernatant and add 5 μl of glycogen (4μg/μl for total 20μg).
6. Add 1000 μl ethanol and centrifuge at 14,000 rpm for 15 min.
7. Wash with 1000 μl ethanol and centrifuge at 14,000 rpm for 10 min.
8. Dissolve in 50 μl EcoR I buffer at 37°C for 15 min and inactivate at 80°C for 20 min. The cccDNA samples are ready for qPCR.

### qPCR of serum HBV DNA and intrahepatic rcDNA and cccDNA

Primers and probe sequences for detection of serum HBV DNA and intracellular rcDNA by qPCR are listed in Table 3 while primers sequences for detection of cccDNA flank the gap region and probe is placed immediately after DR1 sequence (Table 3). The specificity of the listed cccDNA primers and probe can discriminate against rcDNA amplification by 300-6000-fold (data not shown). qPCR was performed using Taqman fast advanced master mix (ThermoFisher cat no:4444558) in a QuantStudio 3 instrument (ThermoFisher cat no: A28136) that accommodates 0.1 ml 96-well hard-shell plate.

**Table 3.**
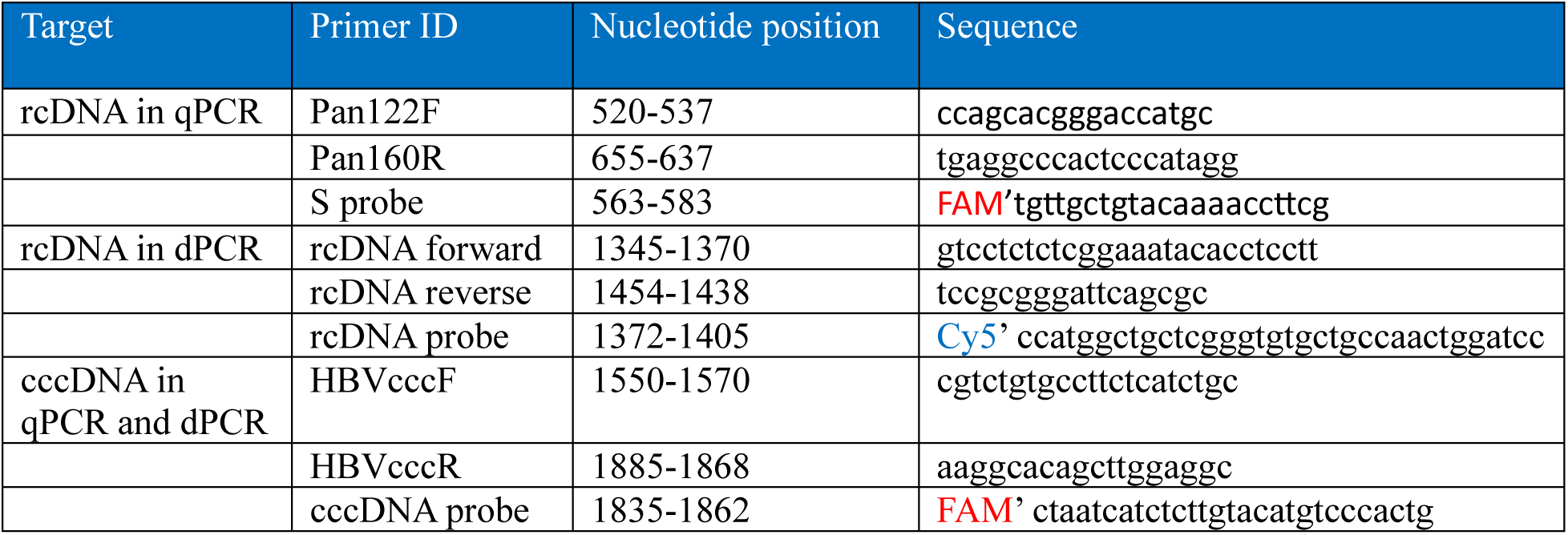
Positions and sequences of cccDNA and rcDNA primers and probes.

All standards used for qPCR were calibrated with the Absolute Q digital PCR.

### The Absolute Q (ABQ) Digital PCR of cccDNA

cccDNA copies/cell were computed initially based on qPCR, then retested with the Absolute Q digital PCR (ThermoFisher cat no: A52864). Briefly, the procedures included the following steps:

1. Prepare 9.1 μl reaction mix consisting of 1.8 μl of 5x DNA dPCR mix (ThermoFisher cat no: A52490), 0.5 μl of 20x primers/probe mix (final concentration 900nM each primer and 250 nM probe), 1 μl cccDNA sample, and 5.8 μl of DNase and RNase free H_2_O
2. Load 9 μl reaction mix into one well of microfluidic array plate (MAP, ThermoFisher cat no: A53301)
3. Run dPCR consisting of 10 min preheat at 96°C and 40 cycles of 5 s at 96°C and 15 s at 60°C
4. Generate data report using QuantStudio Absolute Q Digital PCR software.

The sensitivity of dPCR is a single copy per microchamber and the result deemed valid if Rox fluorescent signal was read in >19,000 of 20,480 microchambers for each sample.

### Principles of simultaneous detection of cccDNA and rcDNA by ABQ duplexing Digital PCR

The ABQ digital PCR instrument can simultaneously detect 4 fluorescent signals of FAM, VIC, ABY, and JUN/Cy5, which allows to detect 4 different targets in the same reaction (Multiplexing). The emission wavelengths of FAM and Cy5 are 517 and 670 nm, respectively, and there is no overlapping in their wavelength spectrum. Thus, FAM was selected for labelling cccDNA probe and Cy5 for labelling rcDNA probe for detection of both molecules in the same reaction, called duplexing dPCR.

The specificity of cccDNA detection is provided through cccDNA specific primers that flank the gap region in HBV genome and the cccDNA specific probe that is placed immediately after DR I sequence (Table 3).

### Linearized cccDNA template cannot generate fluorescent signal with rcDNA primers/probe detection system

HBV DNA released from each of the deposited nuclei will be subjected to NcoI digestion to exclude cccDNA from detection with rcDNA primers/probe. Plus strand in rcDNA is only partially synthesized, containing a single-stranded gap of 600-2100 nucleotides at 3’ end^55^. Since NcoI is located between nt1372 and 1376 that is close to the 3’end of plus strand, it is most likely present in a single strand sequence in rcDNA molecules^55^. Thus, NcoI will linearize cccDNA but cannot cut rcDNA (Figure S2A and B).

The NcoI linearized cccDNA sequence starts with C at nt1373 (5’) and ends with C at nt1372 (3’) (Figure S2C). rcDNA forward primer will bind the 3’ end of the linearized cccDNA, but the rcDNA probe binds its 5’ end. The Taq DNA polymerase that binds the forward primer at 3’ end cannot reach the probe that is located at the 5’ end, thus cannot cut off the 1^st^ base C with Cy5 dye through its 5’-3’ exonuclease activity, and the Cy5 fluorescent signal cannot be generated (Figure S2D). Cy5 fluorescent signal will be generated if both rcDNA forward primer and probe bind a continuous template comprising nt1345 to nt1454 sequentially, that is, F primer binds upstream of the probe binding position (Figure S2C), which only occurs in rcDNA after NcoI cut. Thus, rcDNA, but not cccDNA, will be specifically detected with rcDNA primers/probe following NcoI cut.

We used an HBV DNA plasmid (an ADW subtype monomer cloned into the Psp65 vector) that was linearized by NcoI as a surrogate cccDNA molecule to validate that cccDNA was not detected by the rcDNA probe/primers. The NcoI-digested plasmid was serially diluted and tested using ABQ-duplexing dPCR containing both cccDNA and rcDNA probe/primers. Serially diluted cccDNA molecules were detected using the FAM labelled cccDNA probe; however, no positive signal was detected using the Cy5 labelled rcDNA probe, which detects the extracted rcDNA (Figure S8A and B).

### Simultaneous detection of both cccDNA and rcDNA

The cccDNA samples extracted with the modified Hirt method^56^ contain rcDNA molecules. Deproteinized rcDNA molecules were detected in the extracted cccDNA samples by southern blot^4,47,57^. Therefore, we used the extracted cccDNA samples to evaluate the ability of duplexing dPCR to detect cccDNA and rcDNA. The rcDNA was detected by a regular qPCR in the extracted cccDNA samples using a FAM labelled rcDNA probe (data not shown). Both cccDNA and rcDNA were detected in the extracted cccDNA samples by ABQ-duplexing dPCR (Figure S8C). Our results not only demonstrated the ability of duplex dPCR to detect both cccDNA and rcDNA molecules, but also support the idea that nuclear rcDNA molecules can be used as HBV infection markers.

The sorted nucleus in each well is subjected to NcoI digestion. The DNA sample from a single nucleus will be mixed with dPCR solution containing both ccc and rcDNA primers/probes and loaded into microfluidic array plate (MAP) for dPCR detection. The dPCR results will be generated using QuantStudio absolute Q digital PCR software 6.0.

### Threshold for FAM and Cy5 positive fluorescence

After extensively evaluating fluorescent intensity and distribution pattern among cccDNA positive and uninfected samples, we set the 500 value of both FAM and Cy5 fluorescent intensity as threshold for positive signal. However, we detected about 5% of positive FAM signal and 1% Cy5 positive signal among 576 nuclei prepared from 3 uninfected human livers (2 purchased from PheonixBio and one collected before infection) because there were some free fluorescence molecules remained in each probe despite the standard HLPC based purification of FAM and Cy5 labeled probes. Free fluorescence molecules are not associated with the quencher and can be detected at high intensity without amplification if many free fluorescence molecules are distributed in a microchamber. For example, ROX fluorescence (unlabeled and in free form) is included in ABQ dPCR master mix and distributed to each microchamber for quality control. But the number of ROX molecules distributed to each of 20480 microchambers varies and results in high intensity if many molecules are distributed in one microchamber (Figure S8D). Using the findings from the 3 uninfected livers as reference, we assume that false positive rate for cccDNA and rcDNA by duplexing dPCR were about 5% and 1%, respectively, implying that the true number of cccDNA positive nuclei in each of 3 infected liver samples could be 5% lower than the number detected. But they do not significantly impact the main findings of cccDNA-/rcDNA+ nuclei.

### Detected rcDNA molecules in the nuclei were not non-specifically bound to nuclei

To evaluate the possibility that detected rcDNA in sorted individual nuclei was non-specifically bound to nuclear membrane during preparing nuclei suspension through homogenization that released virions and capsids into lysate, we prepared nuclei suspensions from two livers isolated from two uninfected chimeric mice (animal ID HKB-043-020 or B20 and HKB-043-046 or B46 purchased from PheonixBio). Each nuclei suspension was divided into two vials, one was directly used for sorting, the other was mixed with lysate (containing 0.05% triton-X100) from mouse 987 who was an untreated control with average 870 copies of rcDNA/cell for 20 minutes, then removed the lysate and dissolved in isotonic buffer (154:1 TE with 0.05% triton-X100) for sorting. The sorted nuclei from 4 vials were subjected to duplexing ABQ dPCR. Table 4 shows no significant differences in detecting nuclei with Cy5 intensity ≥500 between two nuclei suspensions mixed with mouse 987 lysate and the two nuclei suspension without mixing, suggesting that detected rcDNA molecules in sorted nuclei unlikely derived from released virions and capsids that non-specifically bound nuclei, which is consistent with the concept that HBV capsid mainly utilizes cellular transport machineries, but not diffusion or passive trapping to reach nuclear membrane where interaction between nuclear localization signal on capsid and nuclear import receptors occurs^58,59^.

**Table 4.** Evaluation of the rcDNA signal in uninfected nuclei with or without mixing mouse 987 liver lysate.

|  | Liver ID: B20 |  | Liver ID: B46 |  |
| --- | --- | --- | --- | --- |
|  | Not mixed with 987 lysate | Mixed with 987 lysate | Not mixed with 987 lysate | Mixed with 987 lysate |
| Number of duplexing dPCR | 96 | 96 | 80 | 96 |
| Number of positive rcDNA signal ($\geq 500$) | 0 | 0 | 2 | 2 |

### Western blot of pre-S and HBsAg in serum and liver lysates

Serum and liver samples were resolved by SDS-PAGE and HBV surface proteins detected by western blot analysis using a rabbit polyclonal anti-HBs antibody (Virostat)^60,61^.

### Immunohistochemical staining of HBsAg on sections

Briefly, formalin-fixed paraffine-embedded liver sections were cut at thickness of 5μM and used for HBsAg staining after deparaffinization, proteinase K digestion, and inactivation of endogenous peroxidase with 3% hydrogen peroxide. Rabbit anti-HBs antibody (LS-C683282, LSBio) and goat anti-rabbit IgG conjugated with HRP (LS-C316062, LSBio) were used as primary and secondary antibody, respectively. DAB chromogen kit (ACH500-IFU, CP Lab Chemicals) was used for color development.

### Statistical analysis

HBsAg (IU/ml) and HBV DNA (copies/ml) and antibody levels (μg/mL) are expressed as mean ± standard deviation (SD). Average intracellular rcDNA and cccDNA levels are expressed as copies/cell. The number of cells per sampling was calculated using sample weight (mg) multiplied by 1.39E5 cells per mg liver tissue^62^, then normalized by factor 0.7 by considering 70% of human liver cells are hepatocytes^63^. The formula to calculate copies/cell is listed below:

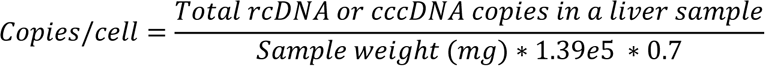

