## Supplemental figure for "Replication-driven HBV cccDNA loss in chimeric mice with humanized livers"

**Supplemental Figure (Figure S) Titles and Legends**

**Figure S1. H&E staining shows unremarkable histology on 2 uninfected and 12 infected liver sections. Blue:** untreated**. Green:** treated**.**

**Figure S2. The NcoI linearized cccDNA template supports the specificity of rcDNA detection. A.** cccDNA is linearized with NcoI**. B.** rcDNA can’t be cut with NcoI. **C.** rcDNA F primer and probe binding positions in cccDNA template linearized by EcoRI**. D.** Showing Cy5 dye labelled 1^st^ 5’ base of the probe can’t be cut by Taq DNA polymerase on the NcoI cleaved cccDNA template**. F primer.** rcDNA forward primer**. Blue arrows:** rcDNA primers**. Blue open arrow:** Cy5 labeled rcDNA probe**. Red arrows:** cccDNA primers**. Red open arrow:** FAM labelled cccDNA probe**.** cccDNA sequence (ADR v00867) is numbered from EcoRI site.

**Figure S3. Sustained high level of anti-HBs antibody expressed by AAV-anti-HBs vector HBV10. A**. The kinetic anti-HBs antibody levels (μg/ml) expressed in both immunocompetent and immunodeficient mice for 252 days post transduction. **B**. Anti-HBs antibody levels (mIU/ml) detected on day 102 pi or earlier among 25 chimeric mice that were administered with HBVZ10 on day 11, 22 or 44pi, respectively. Green, anti-HBs antibody level at 12mIU/ml considered protective among vaccinated subjects. Red, average anti-HBs antibody level among patients with chronic HBV infection, who late cleared HBV infection^1^.

**Figure S4. HBV infection course and intervention schedules among 3 animal experiments. A**. HBV infected mice received anti-HBs antibody (expressed by HBVZ10 and exogenous mouse anti-HBs antibody) monotherapy. **B**. HBV infected mice received anti-HBs antibody (expressed by HBVZ10 then boosted with mouse anti-HBs antibody) and 9-week entecavir therapy. **C**. HBV infected mice received anti-HBs antibody expressed by HBVZ10 (no mouse anti-HBs antibody administered) and 12-week entecavir therapy.

**Figure S5**. **Kinetic serum HBsAg and anti-HBs levels**. **A**. Kinetic serum HBsAg levels. **A1**. Mock treated with AAV vector expressing malaria antibody administered on day 49pi at a dose of 1E11 copies. **A2**. Treated with anti-HBs antibody expressed by HBVZ10 administered on day49pi at a dose of 1E11 copies. **A3**. Treated with mouse anti-HBs antibody started on day 74pi at a dose of 250μg/injection triweekly until day 253pi. **B**. Kinetic serum anti-HBs levels. **B1**. Expressed by HBVZ10. **B2**. Infused with mouse anti-HBs antibody.

**Figure S6. Comparable results of cccDNA quantification between qPCR with ABQ dPCR calibrated standards and ABQ dPCR among 140 cccDNA samples**. **A-G** display cccDNA copies/cell among 20 cccDNA samples from mice 475, 478, 479, 497, 984, 988, and 981, respectively.

**Figure S7. Kinetic serum human albumin levels. A**. Two serials serum samples from experiment 1 were tested at 1/1,000,000 dilution showing higher levels with fluctuations over time. **B**. Seven serial samples from experiment 2 tested at 1/1,000,000 dilution (B1) and two serial samples from experiment 2 tested at 1/100,000 dilution (B2). **C.** Serial samples from experiment 3. Two series tested at 1/1,000,000 dilution (C1). Ten series tested at 1/100,000 dilution (C2). **Blue**: untreated or mock treated. **Green**: treated mice with progressive reduction of serum HBsAg to undetectable level.

**Figure S8. Validation of detection specificity by ABQ duplexing dPCR**. **A**. The rcDNA probe (Cy5) and primers do not detect cccDNA molecules (HBV DNA plasmid linearized with NcoI). **A1**. 100 copies of cccDNA and no rcDNA detected (A1 rcDNA). **A2**. 10 copies of cccDNA and no rcDNA detected (A1 rcDNA). **B**. Cy5 labeled rcDNA probe for duplexing PCR can efficiently detect rcDNA. The same rcDNA sample (about 60 copies/μl) detected FAM labeled rcDNA probe that is routinely used for rcDNA quantification (**B1**) and by Cy5 labeled rcDNA probe for duplexing dPCR (**B2**). **C**. Both cccDNA and rcDNA detected in the same cccDNA sample in the same well by ABQ duplexing dPCR. **C1**. 379 copies of cccDNA detected by cccDNA probe (FAM). **C2**. 3040 copies of rcDNA detected by rcDNA probe (CY5). **D**. showing high ROX intensity signals detected in 3 microchambers while the ROX signals in the remaining chambers were below the threshold.

**Figure S1**

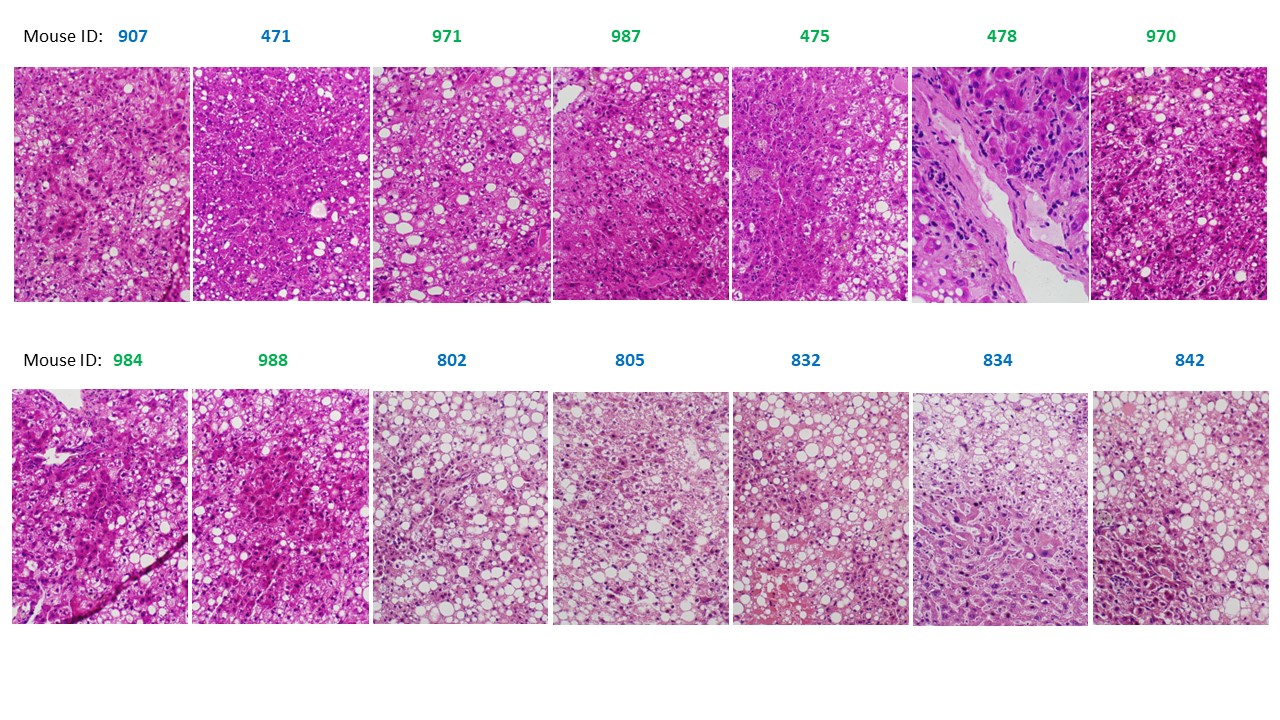

**Figure S2**

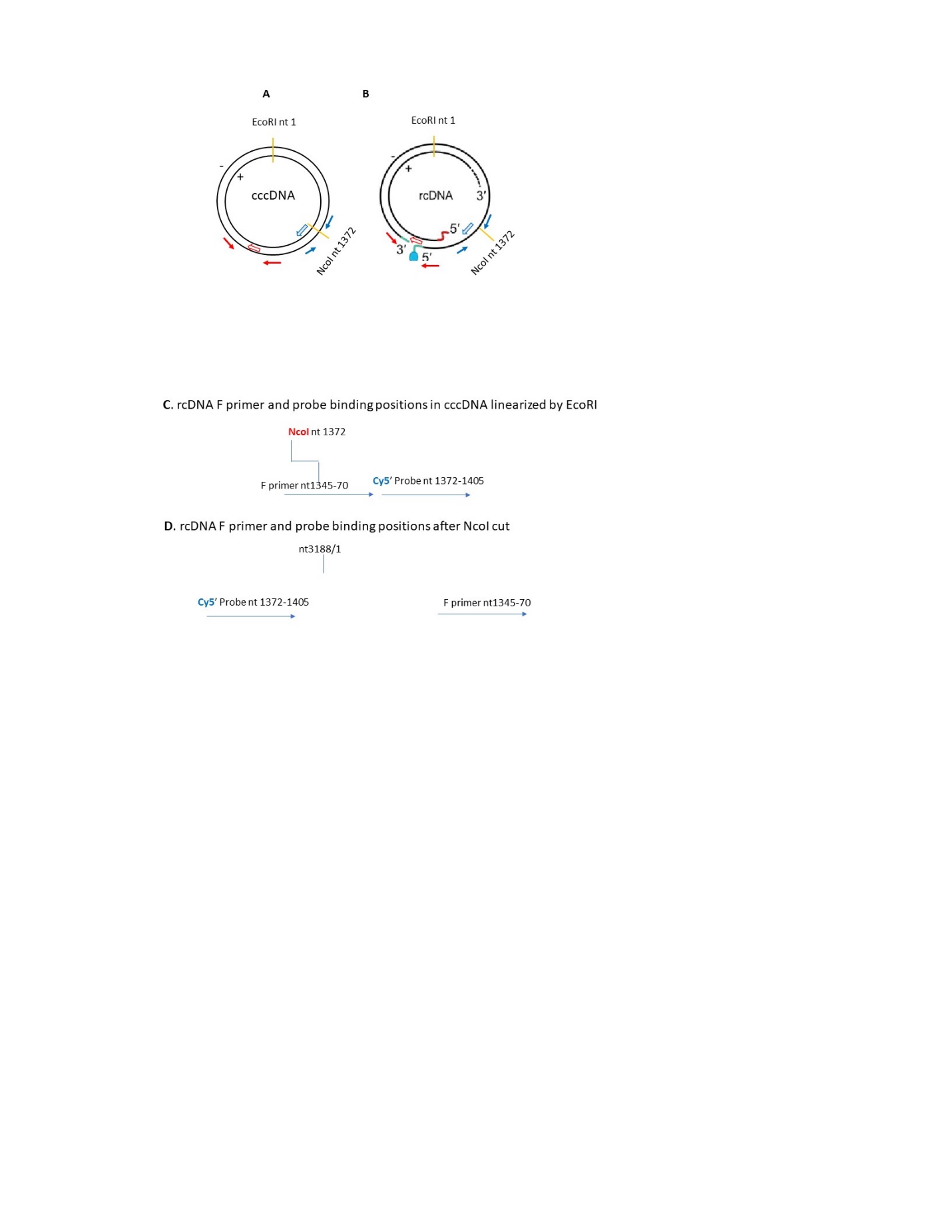

**Figure S3**

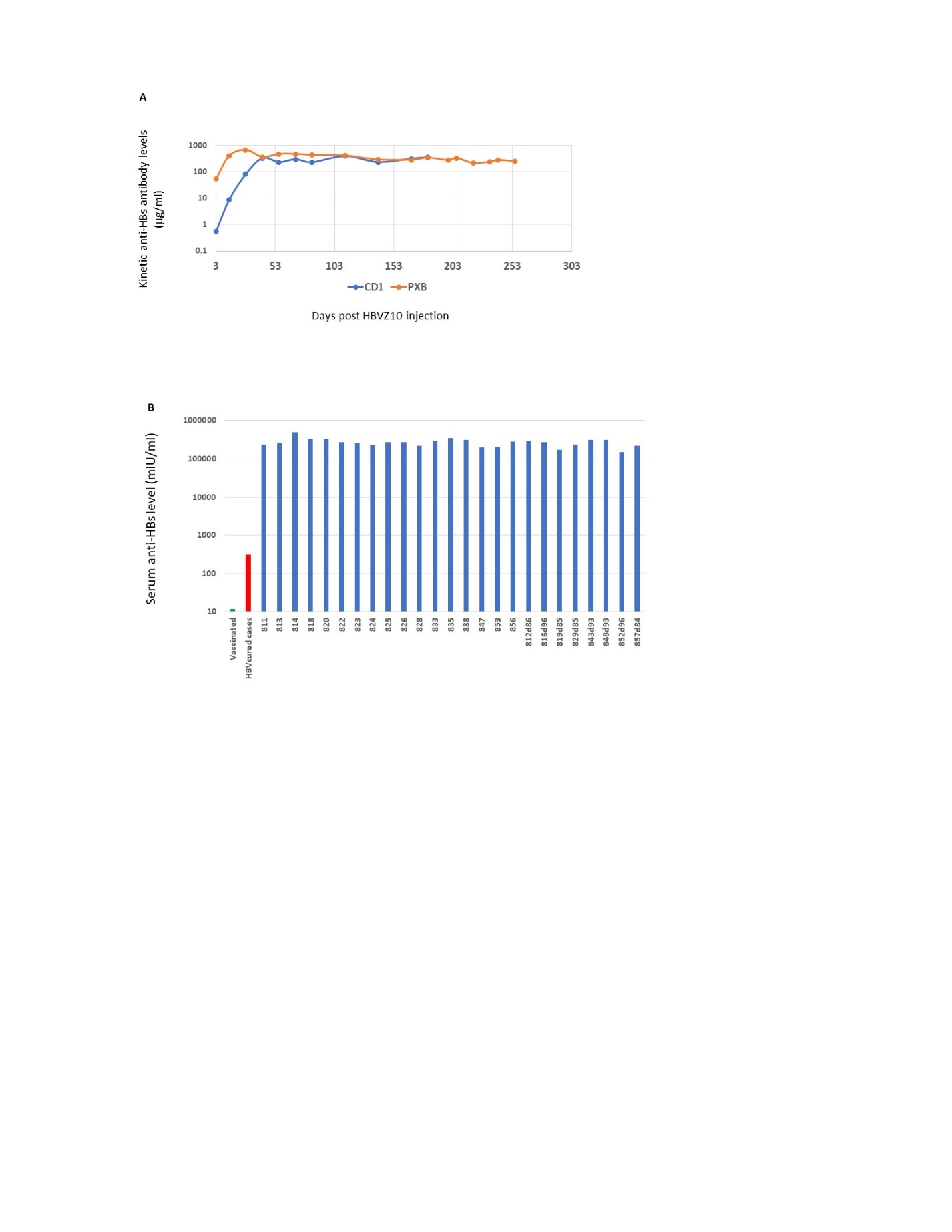

**Figure S4**

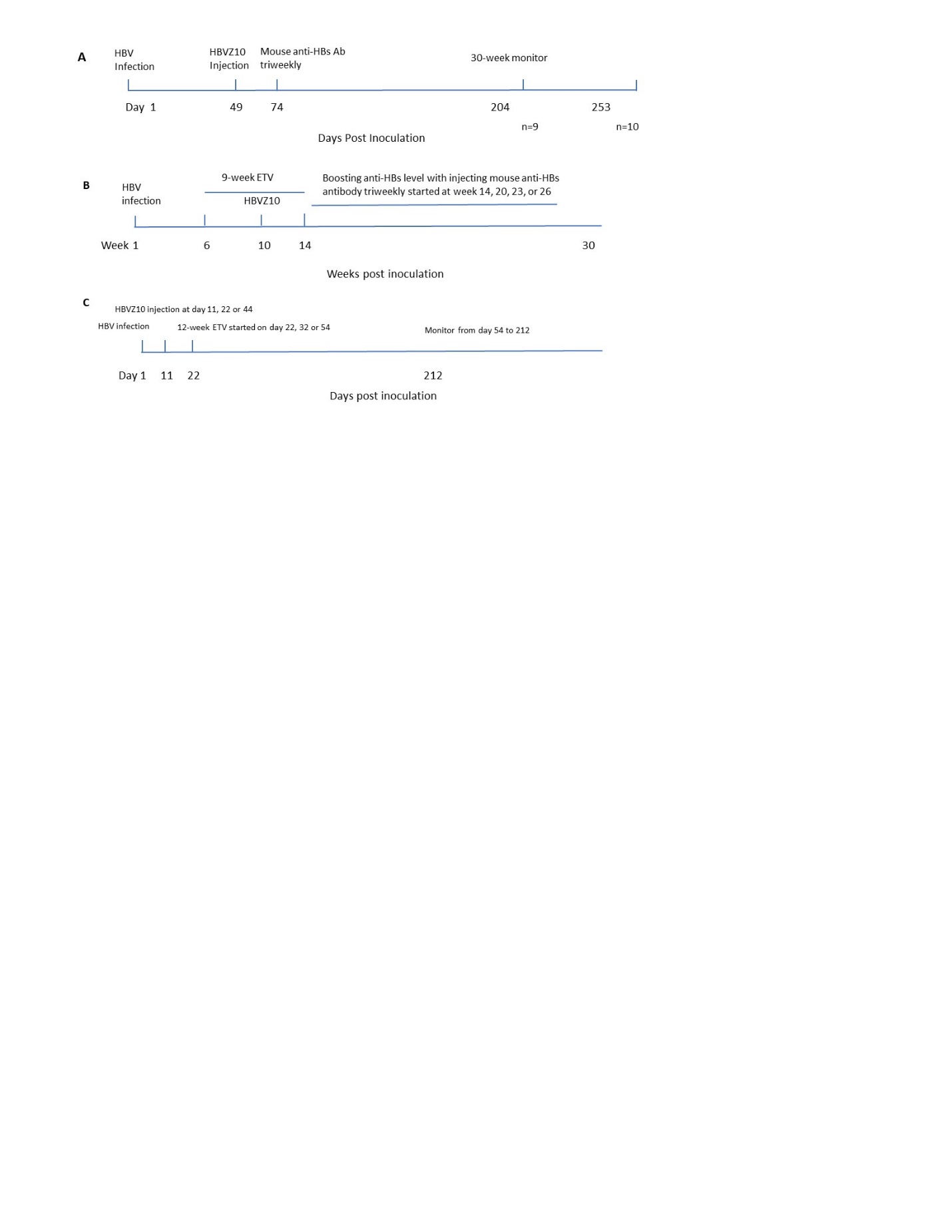

**Figure S5**

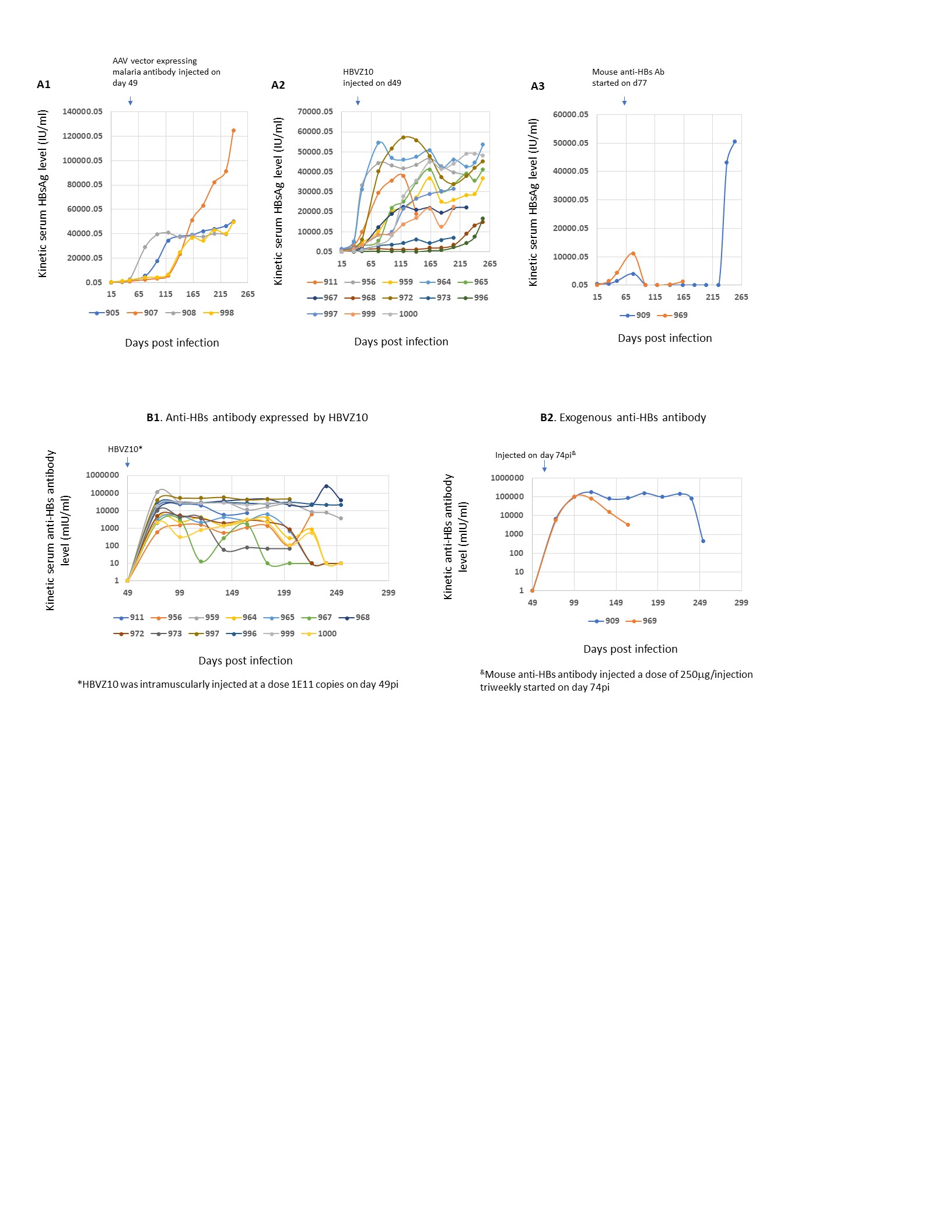

**Figure S6**

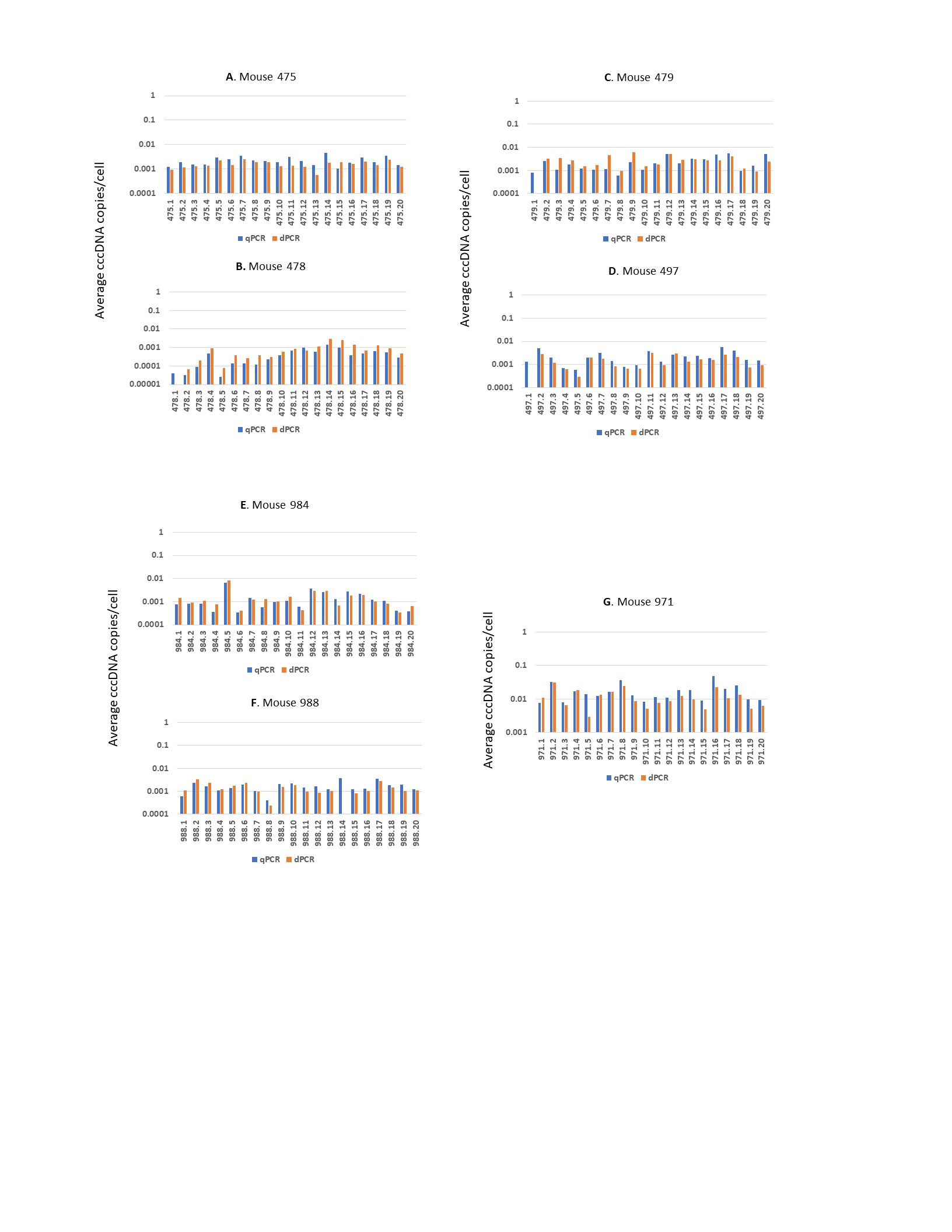

**Figure S7**

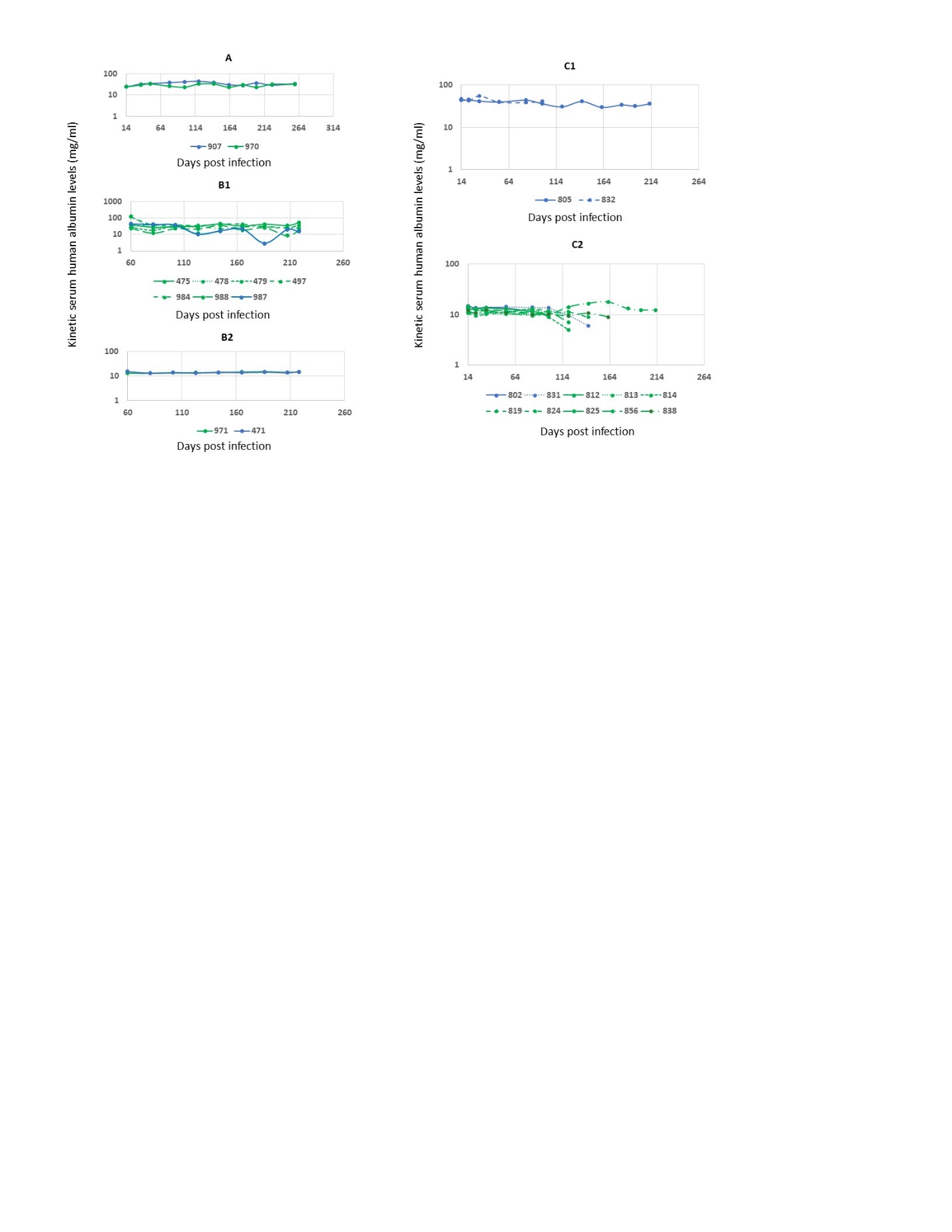

**Figure S8**

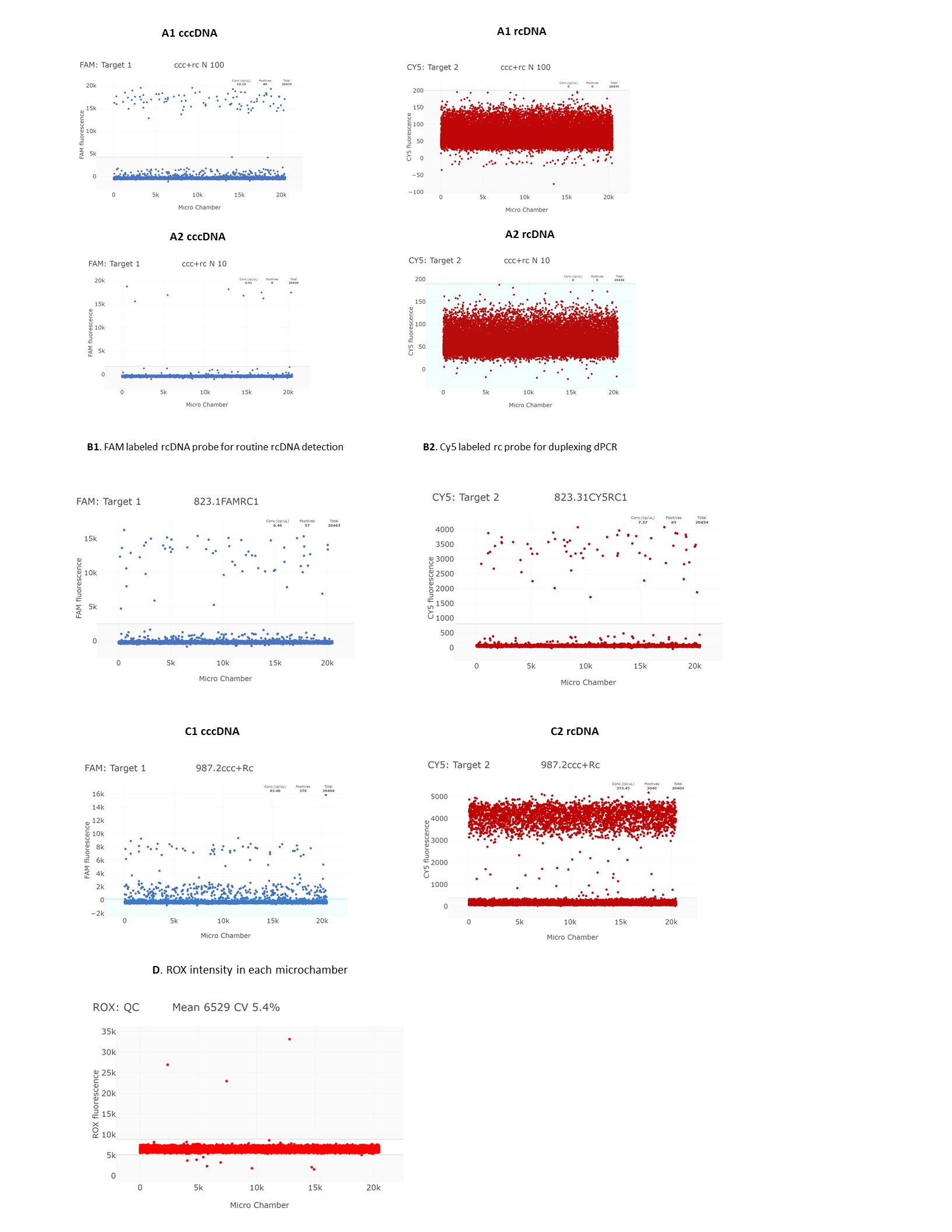
